# Impact of force field polarization on the collective motions of proteins

**DOI:** 10.1101/2024.12.01.626224

**Authors:** Ana Milinski, Annick Dejaegere, Roland H. Stote

**Affiliations:** Institute of Genetics and Molecular and Cellular Biology, 1 Rue Laurent Fries, 67400, Illkirch-Graffenstaden, France

**Author notes:** Corresponding Author: Roland H. Stotes, Institute of Genetics and Molecular and Cellular Biology, 1 Rue Laurent Fries, 67400 Illkirch-Graffenstaden, France.

**Keywords:** Molecular dynamics simulations, nuclear receptor protein, PPARγ, allostery, community network analysis, shortest path method, Drude force field, polarizable force field

## Abstract

Correlated motions of proteins underpin many physiological mechanisms, such as substrate binding, signal transduction, enzymatic activity and allostery. These motions arise from low frequency collective movements of biomolecules and have mostly been studied using molecular dynamics simulations. Here, we present the effects of two different empirical energy force fields used for molecular dynamics simulations on correlated motions – the non-polarizable CHARMM 36m additive force field and the polarizable Drude force field. The study was conducted on two proteins, ubiquitin - a small protein with a well-described dynamic - and the nuclear receptor protein PPARγ. The ligand binding domain of PPARγ was of particular interest since its function is to regulate transcription through ligand and coregulator protein binding. It has been previously shown that a dynamical network of correlated motions ensures the transmission of information related to ligand binding. We present the results of classical MD simulations where we analyze the results in terms of residue fluctuations, correlation maps, community network analysis and Shortest Path Method analysis. We find that the RMS fluctuations tend to be greater and that the correlated motions are less intense when using the Drude force field than when using the non-polarizable all atom additive force field. Our results provide the first quantification of the impact of using a polarizable force field in computational studies that focus on collective motions.

## Introduction

While experimental structure determination has shed light on the many conformations a particular protein can exist in, there remains little in the way of experimental exploration of the detailed motions of particular conformations. Of these, long-range correlated motions are considered fundamentally important for key functional properties of proteins such as substrate binding, allostery and catalysis^1^. Changes in single domain collective motions have been associated to the sensing of ligand binding resulting in the propagation of a signal through the protein to transmit information and alter activity. Studies have suggested that correlated motions of secondary structure elements, such as β-sheets, contribute importantly to protein function ^2^. For example, PDZ domains are protein interaction modules that recognize short amino acid motifs at the C-termini of target proteins. Ligand binding affects the transfer of binding information to other domains in the context of PDZ-containing multidomain scaffold proteins. In the PDZ domain, the global network of correlated motions, can lead to the coupling of the N- and C-terminal ends by pathways involving the β-sheets. These motions arise from the low-frequency collective movements of residues and it has been suggested that these protein motions are selected by evolution^3,4^.

While the importance of correlated motions has become more apparent and appreciated, they remain difficult to measure experimentally, so one of the principal methods for studying correlated motions is by molecular dynamics simulations. Molecular dynamics simulations of proteins rely on the use of empirical force fields, which are parameterized using, for the most part, experimental data and quantum mechanical calculations. And while this approach has been used with great success over the past decades to study a wide range of topics, there is a constant effort to introduce improvements. One such effort has been to improve the treatment of electrostatic interactions, which in standard classical force fields, are treated by fixed point charges. Efforts by numerous teams have focused on introducing aspects of electronic polarization. One approach characterizes the charge redistribution within each atom, by either induced dipoles ^5^ or by a Drude oscillator model ^6^, and the other approach is based on charge flow between atoms, as implemented in the fluctuating charge (FQ) model ^7^.

Developed largely in the framework of the CHARMM all-atom force field, the Drude oscillator model for protein force fields ^8^ is a theoretical framework that introduces an auxiliary particle called the “Drude particle” for each atom representing a loosely bound electron that contributes to the atomic polarizability. A harmonic oscillator function is used to connect the Drude particle to the atom, simulating the restoring force on the electrons. The model introduces a term that represents the polarizability of the atom, allowing for the simulation of electronic response to an external electric field. The development of the Drude model in the context of polarizable force fields in molecular dynamics simulations aims to capture electronic polarization effects that are not explicitly represented in the classical models. This type of model is intended for studying systems where electronic polarization plays a significant role.

The Drude model has been used and benchmarked for a variety of systems ^9^, and several reviews are available ^10,11^. Though the Drude model for polarization has undergone extensive development and application, the analysis associated with the applications has largely focused on aspects of structure, energetics and local dynamics. None of these studies have utilized the Drude model to address questions concerning collective behavior.

Collective motions occur across the three-dimensional structure of the protein and the principal tool for studying such motions in proteins is molecular dynamics simulations. The question naturally arises, then, to what degree does the inclusion of polarization affect the collective motions of single protein domains. In this work, we address this question through the study of two proteins, ubiquitin, which contains 76 residues, and the ligand binding domain of the nuclear receptor PPARγ, which contains 276 residues. We assess the impact of the polarization on various dynamical properties, including their collective motions.

## Ubiquitin

Ubiquitin is a small protein that plays a crucial role in various cellular processes, primarily as a regulator of protein degradation. It is found in nearly all eukaryotic cells and is highly conserved across species. Consisting of 76 amino acids, it has a highly conserved three-dimensional structure, with a characteristic beta-grasp fold (Fig. 1).

**Figure 1:**
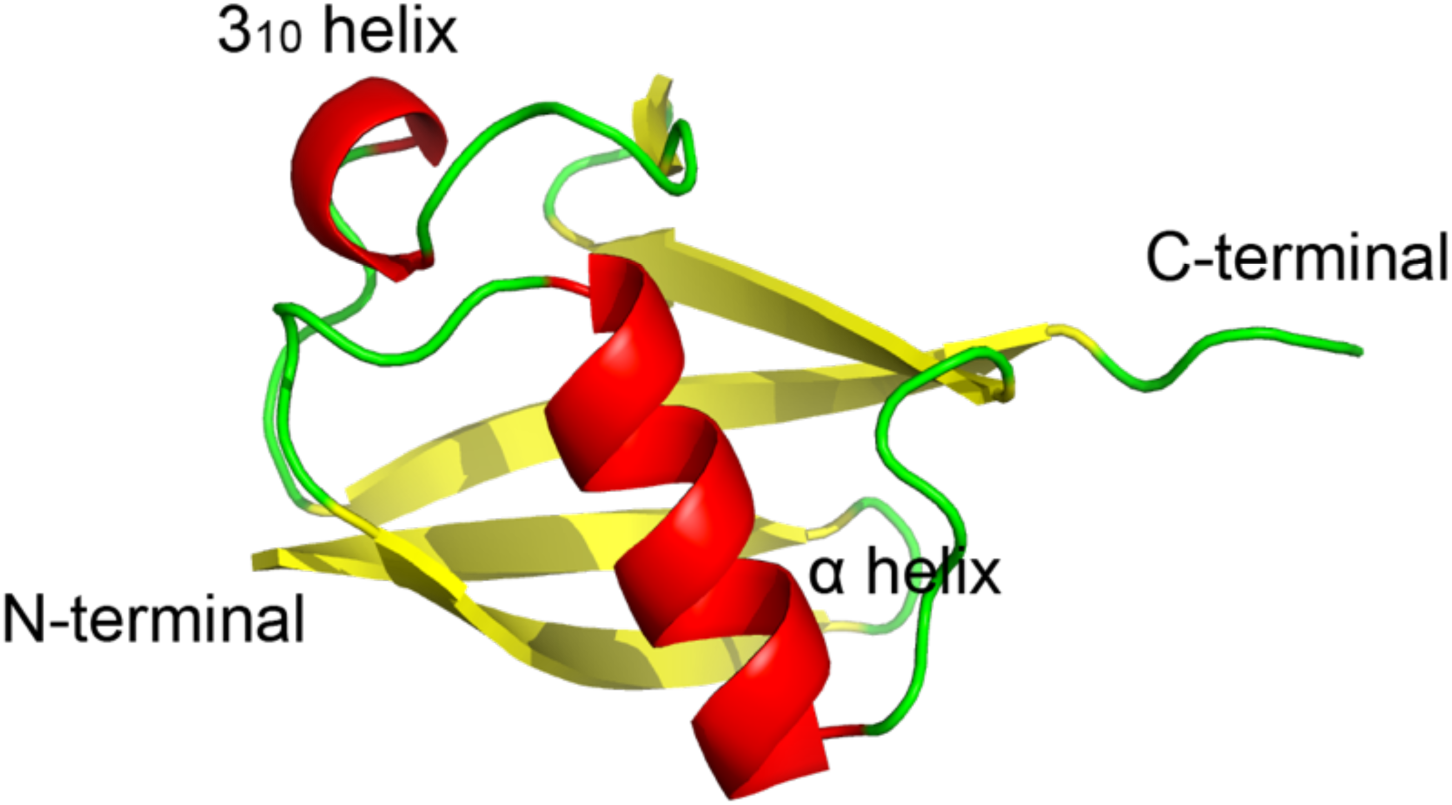
The 3D structure of ubiquitin (1.8Å resolution, PDBID 1UBQ). Shown is the mixed parallel-antiparallel β sheet in yellow, the alpha helices in red and loop regions in green.

A beta-grasp consists of a 4 stranded anti-parallel sheet and a single helical region with a β(2)-α-β(2) topology ^12^. The protein has a flexible C-terminal tail, which is involved in the attachment of ubiquitin to target proteins. Ubiquitination is involved in the regulation of various cellular processes, including cell cycle progression, DNA repair, signal transduction, and immune responses. Its primary function is to mark proteins for degradation by the proteasome. Collective motions in ubiquitin have been suggested to play a role in a conformational switch^13^. Furthermore, a correlation network in ubiquitin was identified as spanning the β-strands ^14^.

In another study, an allosteric switch governed by a collective motion that affects protein– protein binding was extensively characterized and validated using a combination of techniques, including NMR, X-ray crystallography, computer simulation, and enzyme inhibitor assays. These studies suggested that the loops are involved in a pincer-like movement involving residues in the loop between the first (β1) and second β-strands (β2) ^15,16^.

### Nuclear receptor PPARγ

Peroxisome proliferator-activated receptor gamma (PPARγ) is a ligand-dependent transcription factor belonging to the nuclear receptors superfamily ^17^. PPARγ has the common nuclear receptor organization of five conserved domains, including DNA binding (DBD) and ligand binding domain (LBD). Upon signaling events, modifications in structure and/or dynamics relay signaling events to downstream effectors, in particular coregulator proteins. The natural agonist ligands of PPARγ are polyunsaturated fatty acids (PUFAs) and their derivatives ^18^. Its implication in various diseases, such as obesity, cardiovascular disease and diabetes mellitus, makes its ligand binding domain (LBD) an important pharmacological target. Various synthetic agonist and antagonist ligands have been shown to regulate PPARγ activity, notably of the family of glitazones ^19^. Structures of PPARγ LBD in its apo and corepressor-bound form in complex with a peptide from the NCor1 corepressor protein are shown in Fig. 2, along with a cartoon drawing labeling secondary structure elements.

**Figure 2.**
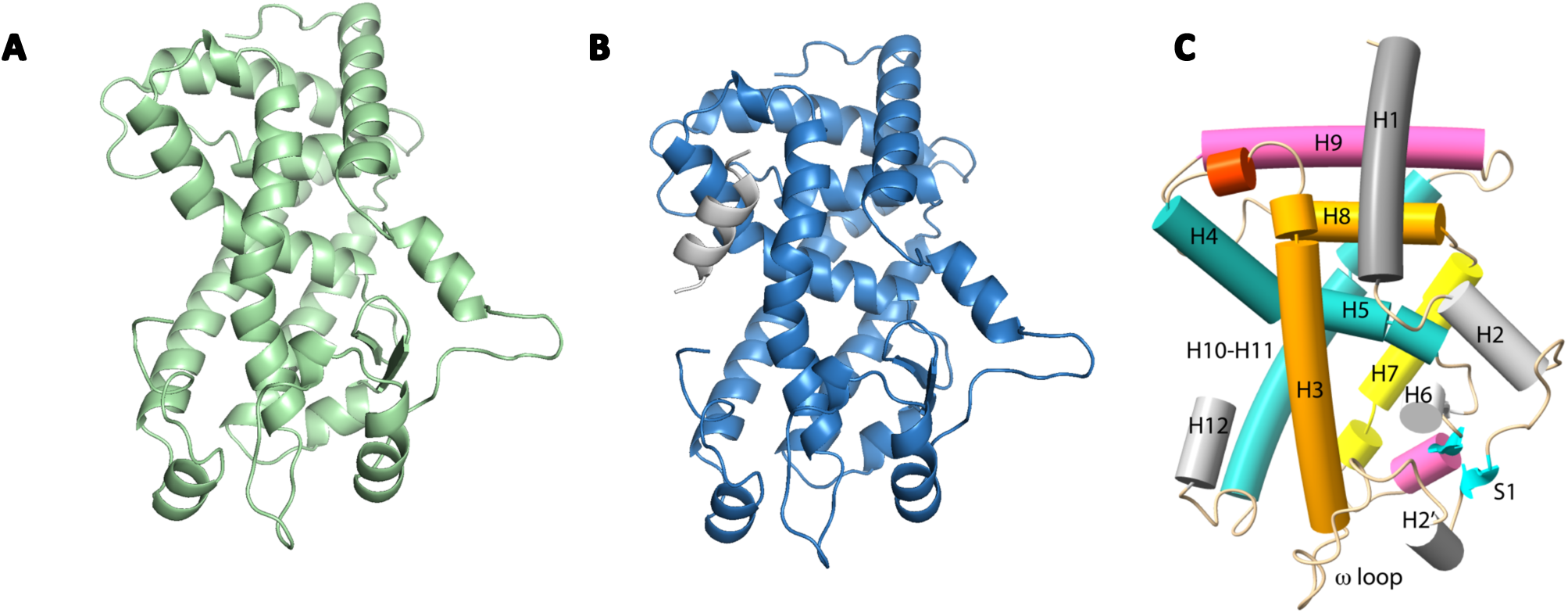
PPARγ ligand binding domain (residues 230 – 505) from the 3.2Å crystal structure PDBID 7WOX chain B, A. in apo form (green) and B. the same LBD modeled (blue) with the corepressor-peptide (gray). C. The secondary structure elements are labeled from helix 1 to helix 12.

The physiological function of NRs is highly dependent on conformation and structural dynamics modulated by ligand binding. The ligand binding domain acts as a dynamic hub, transmitting binding events to other protein interfaces and domains. The PPARγ activation involves a conformational change of the H12 helix at the C-terminal part of the LBD induced by ligand binding^20^. Helix H12 undergoes a transition from a flexible ensemble of conformations to a folded conformation localized on the core of the LBD. Characterized by numerous crystallographic structures of agonist-bound LBDs, this position of the H12 helix is often referred to as the transcriptionally active conformation ^21,22^. In this active conformation, H12, along with helices H3 and H4, constitute a hydrophobic interface called an activation function 2 (AF2). This interface serves as a platform for coactivator protein binding and the recruitment of chromatin modulator complexes as well as other components of basal transcriptional machinery ^23^. In contrast to the active conformation of the LBD and H12, the inactive conformation of the receptor is not structurally well described. The crystallographic and computational data suggest an ensemble of conformations for H12, meaning that this region, in the absence of an agonist ligand, is flexible ^24^.

The first study of the functional dynamics of the PPARγ ligand binding domain (LBD) was done by Fidelak et al ^25^. This study explored the role of allostery in the functioning of the receptor by comparing the LBD in apo and in agonist-bound forms. A dynamical pathway linking amino acids that are in topological proximity and at distance was established, explaining correlated motions primarily arising from low-frequency collective motions. The analysis of correlated motions showed coupling between distant regions of the LBD, such as different helices, the N- and C-termii and other physiologically relevant interfaces, such as the co-regulator binding surface and the dimer interface by which PPARγ interacts with its partner NR, RXRα. As a consequence, changes in this network could impact the ability of LBD to bind ligands and coregulators, and by extension the overall function of PPARγ. Correlated motion calculations and network analysis were later done for the full PPARγ/RXRα heterodimer structure in complex with DNA ^26^.. The results showed the existence of longer range interdomain correlations which were used toward the understanding of allostery in nuclear receptor complexes ^26^. Here, we explore the intrinsic dynamics of PPARγ LBD in its apo- and corepressor peptide bound forms using both the classical all-atom additive empirical energy functions and the Drude force field. We provide quantitative insights into the effects of polarization on the modeling of correlated motions of PPARγ.

## Methods and Analysis

The two systems studied were human ubiquitin and the ligand binding domain (LBD) of human PPARγ. Human ubiquitin has been previously studied by molecular dynamics simulations using the Drude polarizable force field in the benchmark work of Kognole et al^27^. Here, we examine that system from the point of view of collective motions. Secondly, we study the ligand binding domain (LBD) of human PPARγ because of the importance of collective motions in its function. PPARγ is also a larger protein than ubiquitin with a different fold. Each system was prepared using the PDB Reader and Manipulator option of the CHARMM-GUI web interface ^28^ to prepare the simulations using the CHARMM all-atom additive force field (AA) using the CHARMM36 all-atom parameter set ^29^. The CHARMM-GUI Drude Prepper interface ^27^ was subsequently used to prepare the systems for simulations using the Drude polarizable force field.

For ubiquitin, coordinates were obtained from the 1.8 Å resolution crystal structure (PDB ID 1UBQ) ^30^. The protein was solvated in a 64 Å cubic box with 7,921 TIP3P water molecules. The protein carried no net charge and simulations were performed without any neutralizing counterions or added salt. Harmonic constraints were put on the protein and the protein was subjected to an energy minimization followed by a 400ps equilibration simulation. The constraints were removed and the system was run for 400ns. The molecular dynamics simulations were done using the NAMD program under NPT conditions ^31^. For the simulations of human ubiquitin using the Drude force field, the protein was solvated in a cubic box sized 64 × 64 × 64 Å^3^ with 7,921 “simple water model with four sites and negative Drude polarizability” (SWM4-NDP) model ^32^. The structure was subjected to an energy minimization followed by a 100ps equilibration simulation using a 0.5fs time step. The shorter than conventional time step was used to avoid instabilities in the simulations using the Drude force field. The system was then run under NPT conditions for 200 ns using a 1fs time step. The particle mesh Ewald method was used to treat the electrostatic interactions.

For the PPARγ ligand-binding domain (residues 230 - 505), we used the 3.2 Å resolution crystal structure of chain B from the PDB file 7WOX ^33^. Although one chain in this PDB entry is bound to the antagonist MMT-160, the second chain (chain B) did not show any electron density representing a ligand in the binding pocket, so it was taken to be a structure of the apo protein. The protonation states of the histidine residues of this chain were determined using PROPKA method ^34,35^ via the webserver https://server.poissonboltzmann.org/pdb2pqr followed by manual verification and the structure was further prepared using the CHARMM GUI interface ^28^.

The molecular dynamics simulations were done using the NAMD program under NPT conditions^31^. The protocol consists of four steps, first, the protein was fixed, but the water and ions were without constraints. The system was subjected to 1000 steps of energy minimization to allow the water and ions to adjust position in response to the presence of the protein. Next, the system was heated up to 600K, during 23000 steps, again with the protein fixed. This was followed by another energy minimization for 1000 steps. This was followed by a heating to 296.5 K. The constraints on the protein/ligand were removed and the entire system was energy minimized for 2000 steps. The entire system was then heated up to 296.5K over 15000 steps, followed by an equilibration run of 85 000 steps of dynamics then followed by the production phase. A 1 fs time step was used. The duration of each simulation was 100 stages of 1×10^6^ timesteps, which resulted in 100 ns - long simulations. The last trajectory frame was taken as a starting structure for building the structures for the Drude and AA simulations used in this study.

The two systems we compared were the apo form of PPARγ LBD and the PPARγ LBD complexed to the co-repressor peptide NCoR ID1 (12 amino acid sequence GLEDIIRKALMG), identified here as the corepressor-bound form. The coordinates for the corepressor peptide were taken from the crystallographic structure of a PPARγ mutant complexed to the NCoR peptide resolved by our team, referred to here as the in-house structure (unpublished data). The 12 amino acid peptide was added by superposing the apo PPARγ LBD structure on the in-house structure.

For the simulations with the Drude force field, the apo- and corepressor-bound structures were solvated in 100Å cubic water box using the SWM4-NDP water model. A minimization for 2000 steps was done followed by an equilibration for 200000 steps using the NAMD program with the time step of 0.5 fs. During the production phase, we used a time step of 1fs. The duration of each simulation was 100 ns. Three replica simulations were carried out for the four PPARγ LBD systems. Similarly, three replica AA simulations were performed.

For each simulation, the root-mean-square coordinate difference (RMSD) and residue averaged backbone atomic root mean square fluctuations (RMSf) were calculated. The calculated fluctuations were compared to the atomic fluctuations calculated from experimental B-factors. Cross-correlation coefficients were calculated from the molecular dynamic simulations following the equation:

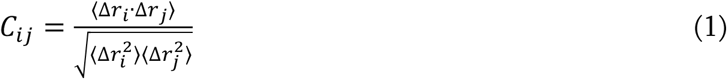

where *r_i_* and *r_j_* are the displacements from the mean position of residues i and j, respectively. From the C_ij_ correlation coefficients, which are organized as a matrix, a cross-correlation map was calculated using a color-coded 2D representation. In this representation, C_ij_ = 1 identifies correlated motions and *C_ij_* = −1 anti-correlated motions. These values give us information concerning the global collective motions. The *C_ij_* correlation coefficients were evaluated over 2ns segments of the simulations. For each segment, a mean structure was calculated and the *C_ij_* correlation coefficients were calculated for the backbone atoms. Correlation maps were obtained by averaging the *C_ij_* over all time interval blocks corresponding to the time interval. Community network analysis was performed using the Bio3D software ^36^. Contact maps were produced using an atom-atom distance cut-off of ≤10 Å and the correlated motions were obtained from the molecular dynamics simulations. The Girvan–Newman algorithm ^37^ as implemented in Bio3D was then used for the community detection. The Girvan–Newman method is a graph-based network approach that is based on the edge-betweenness centrality measure, where the edge-betweenness centrality of an individual residue is defined as the number of the shortest paths connecting other residue pairs that pass through it along the MD trajectory, thus providing an estimate of the influence of this residue on communication, or modularity. Communities of residues are characterized by high modularity values, that is, residues in the same community share dense connections, whereas residues of different communities have sparse or no connections at all. The size of a node is related to the size of a community and a larger sphere depicts a higher number of residues in the node. The edges connect coupled communities, where thicker edges correspond to higher degree of correlation. The correlation threshold for edge detection (*C_ij_* cutoff) was 0.5. The community map analysis results are depicted using colored spheres mapped on the average 3D structure in tube representation.

The Shortest Path Method (SPM) was used to assess the importance of individual residues and their pairwise connections in the structural dynamics of proteins ^38^. This is in contrast to the community network analysis, which establishes communities around multiple residues.

The SPM method produces a network graph based on mean distances and correlation values, and computes shortest path lengths using Dijkstra algorithm ^39^. The shortest path is the most direct path with the most significant connection between two residues and shows how the residues are connected by the structural dynamics of the protein. The tool is mostly aimed at exploring key residues implicated in enzymatic activity, but here we use as a way to assess the similarities and differences of simulations using different force fields. The online tool provided at https://spmosuna.com was used with default values.

## Results

### Structural dynamics of Ubiquitin

#### Drude FF simulation shows larger RMSD time series values

For the simulations of ubiquitin, the backbone RMSD time series was calculated over the production phase of the simulations using the initial structure as the reference structure. For each frame in the trajectory, the complexes were reoriented over the entire backbone. Ubiquitin simulated both with the AA force field and with the Drude force field show a stable RMSD during the trajectories (200ns for the simulation with Drude and 400ns for the simulation without), see Fig. 3. The RMSD values reached plateau values of 1.8 and 2.8 Å for the AA and Drude simulations respectively, so the Drude simulation yields a slightly higher RMSD than the AA simulation. No large-scale conformations were observed during either simulation. The 2.8Å plateau value for the RMSD in the Drude simulation is very similar to what was observed in the work of Kognole et al ^27^.

**Figure 3:**
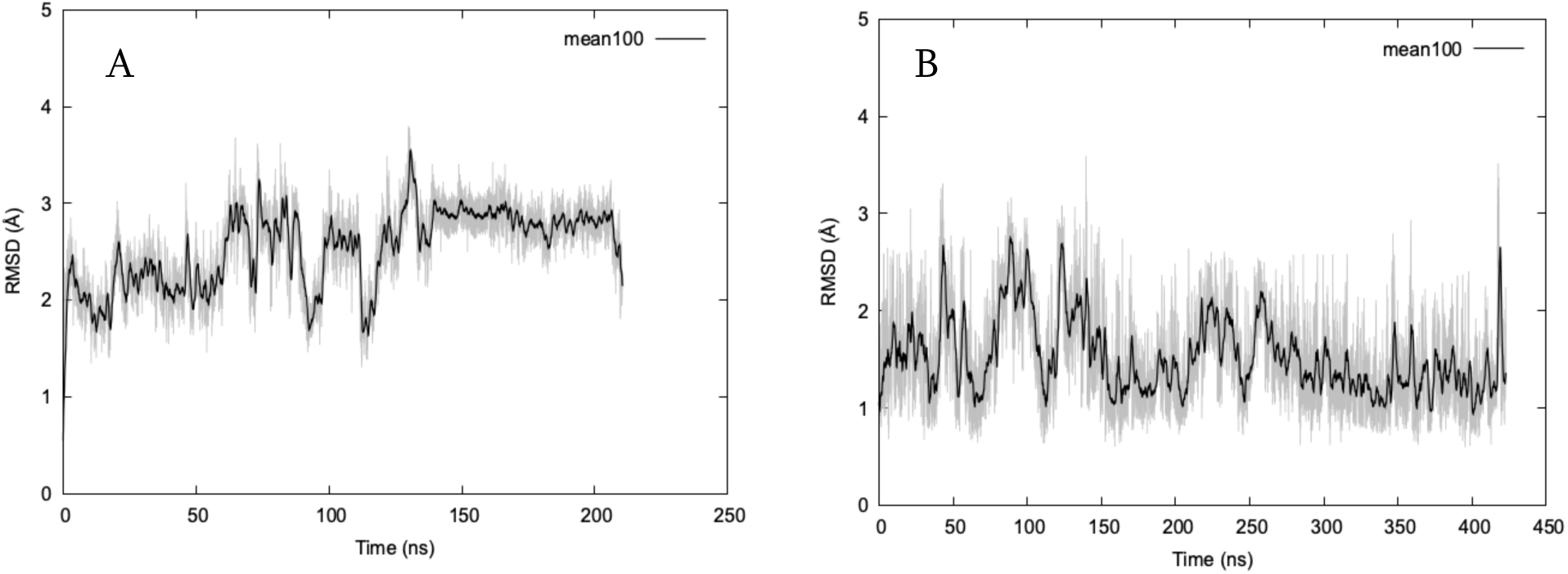
RMSD of ubiquitin backbone atoms with respect to the initial structure. Results from the simulation using the AA force field (A) and the simulation using the Drude force field (B) are shown in gray, black lines correspond to the running average over 100 data points.

#### Drude force field simulations show more significant RMS fluctuations

The by-residue root mean square fluctuations of the backbone atoms for ubiquitin were obtained from the AA and Drude force field simulations (Fig. 4). The fluctuations are in good agreement along the entire protein sequence, although the fluctuations from the Drude simulation are more pronounced than those calculated from the simulation using the AA force field. Although the loops between the α-helixes show more significant flexibility, the fluctuations of the residues within the α-helixes are lower in amplitude. The loop fluctuations tend to be greater in the Drude simulations, but aside from the increases in the loop regions, the fluctuations are very similar in both simulations. The profiles obtained compare well to the profiles presented by Kognole et al ^27^.

**Figure 4:**
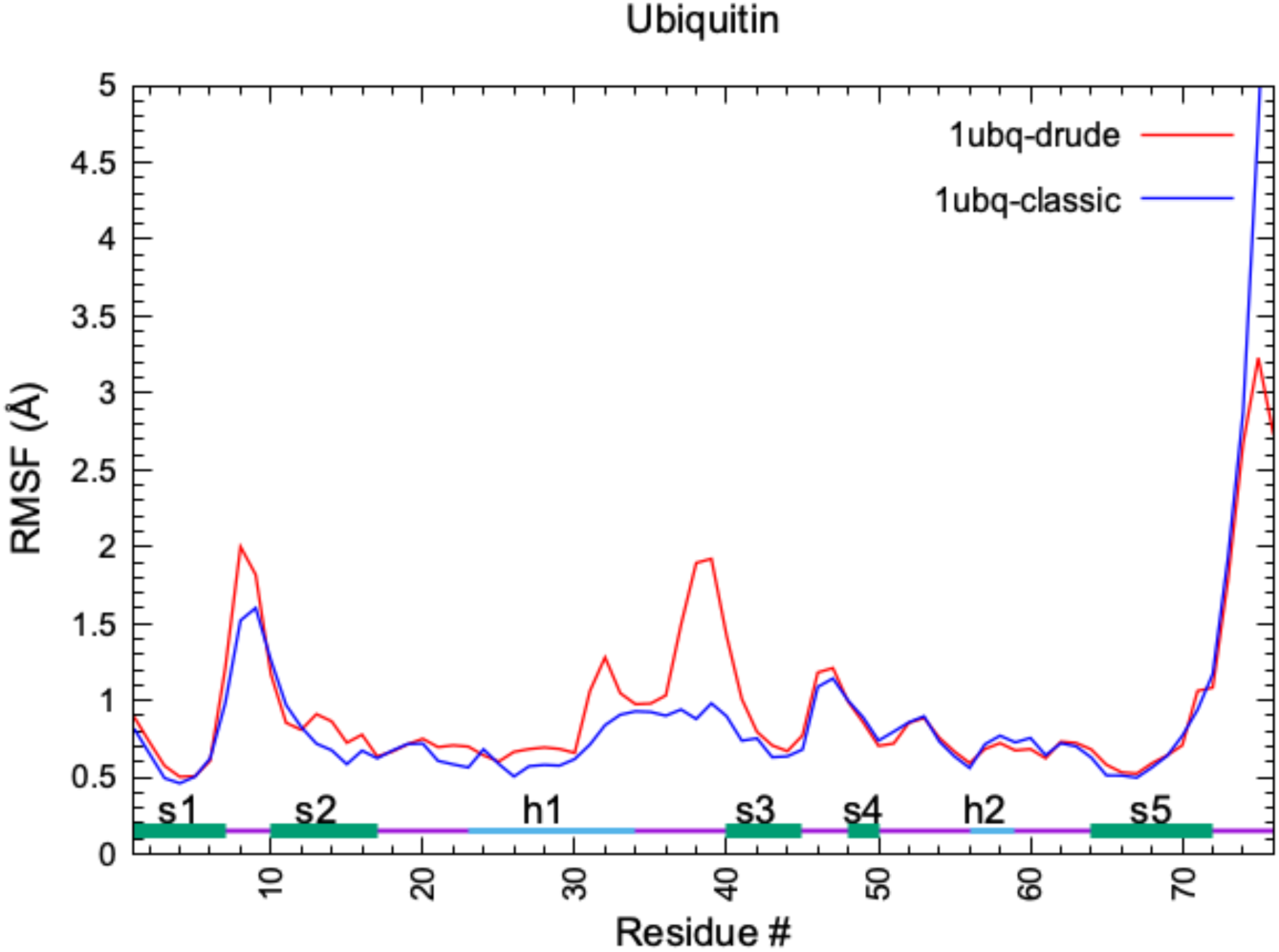
RMSF of backbone atoms averaged by residue for the drude simulations (red), the AA simulation (blue) and from the crystallographic B factors (black). The secondary structure elements are indicated.

Additionally, we plotted the RMS fluctuations of all heavy atoms averaged as a function of the distance from the center of geometry. Related to the Debye-Waller factor, the radially averaged fluctuations provide an overall measure of the motion within the protein. We see from Fig. S1 that the overall flexibility increases as one moves toward the surface of the protein and the behavior is very similar for both the AA and Drude simulations. The AA simulation has a slightly more extended C-terminal end which exhibits higher fluctuations. In both cases, the radius of gyration remains essentially equivalent at 11.7 and 11.8 Å for AA and Drude simulations respectively.

### Structural dynamics of PPARγ

#### Drude FF simulation shows larger RMSD time series values

The RMSD time series for PPARγ were calculated from the molecular dynamics simulations for the three replicas of the AA and Drude simulations for each system. For each system, the average times series of the three replicas were displayed along with the high/low values at each time point. All four PPARγ systems show stable 100ns trajectories (Fig. 5).

**Figure 5.**
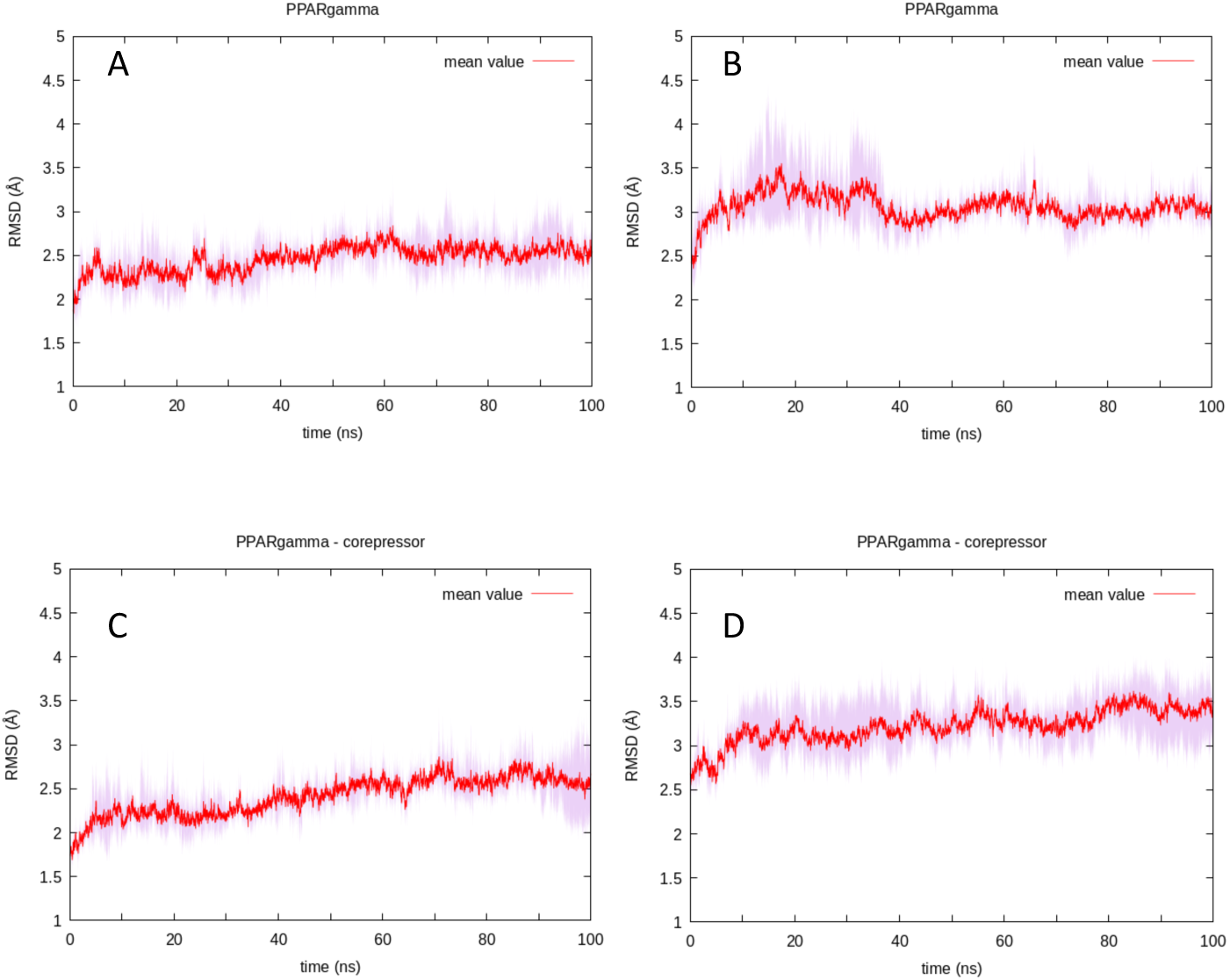
RMSD of PPARγ LBD by AA simulation (A), PPARγ LBD by Drude simulation (B). PPARγ LBD bound to the corepressor peptide by AA simulation (C); PPARγ LBD bound to the corepressor peptide by Drude simulation (D). The mean value of 3 replicas is represented as a red line.

The RMSD mean value of PPARγ apo system simulated with the Drude model was higher than the value of the system simulated with the AA force field with the values being 3 Å (std_dev: 0.09) and 2.5 Å (std_dev: 0.06), respectively (Fig. 5 A,B).

We notice the same trend when comparing the simulations of PPARγ bound to the corepressor peptide NCoR (Fig. C,D). The Drude simulations presented higher values of RMSD, with the mean value of 3.2 Å (std_dev: 0.21) than the non-drude simulations, where the mean value is 2.4 Å (std_dev: 0.04). These results are consistent with the conclusions that the Drude force field allows for a higher conformational flexibility than the standard additive CHARMM force field ^6^. In addition to the overall stability of the PPARγ - corepressor bound complex, we see the interaction of two components as stable as the peptide does not dissociate from the PPARγ LBD. In our case, the Drude force field maintains the protein – peptide complex.

#### Drude force field simulations show more significant RMS fluctuations

We calculated the RMSF of the backbone atoms of PPARγ averaged by residue over all three replicas. In all of the cases, we observe an RMSF profile that well reflects the structure, in that loops are more flexible than the secondary structure regions. Where the AA simulations present the highest flexibility in the regions of the loop between H2 – S1, loop H9 - H10 and the H12 (Fig. 6A,C), the Drude simulation shows a higher flexibility of the Ω loop region and smaller flexibility of the loop H9 - H10 (Fig. 6B,D). The explanation for this could be that in both Drude simulations, with and without corepressor peptide, a salt bridge between residues D411 (H8) and H453 (loop H9-10) is maintained. This salt bridge is present in the crystal structure, and so in the initial simulation structure, and was assigned to be a protonated histidine. This salt bridge is known to be conserved in all NRs of class II ^40^, so the simulations with the Drude force field lead to the maintaining of this conserved salt bridge, while it was lost in the AA simulations. The second salt bridge of the class II NRs, between residues E352 (mid H4 – 5) and R425 (loop H8 – 9), is well maintained in both AA and Drude simulations. Time series for the salt bridges are given in Supplementary Material, Figs. S5A and S5B, for apo PPARγ and PPARγ complexed with the corepressor peptide, respectively. The RMSF variability, represented by the light pink regions, of three replicas is also higher for Drude simulations.

**Figure 6.**
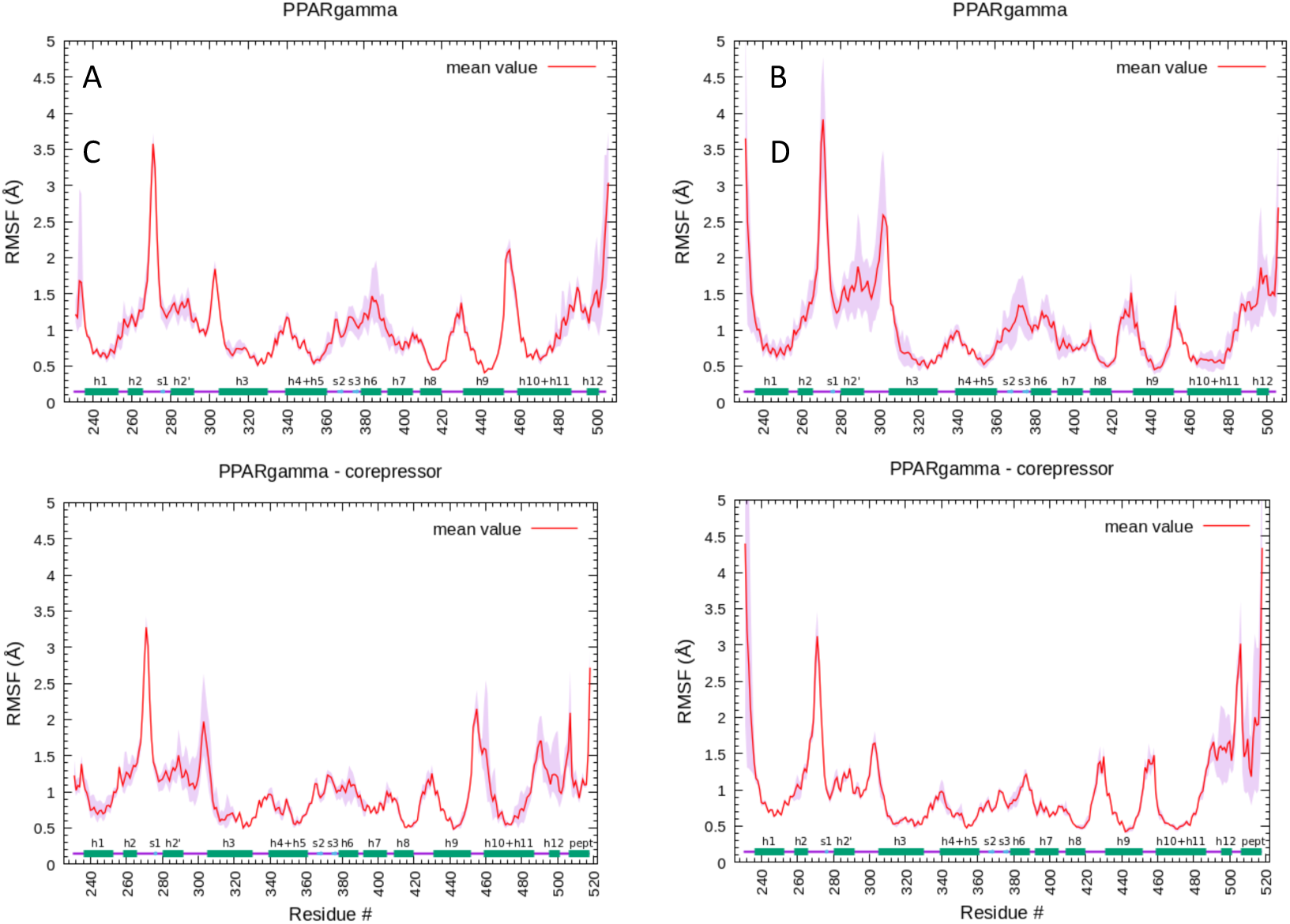
RMSF of PPARγ LBD by by AA simulation (A), PPARγ LBD by Drude simulation (B). PPARγ LBD bound to the corepressor peptide by AA simulation (C); PPARγ LBD bound to the corepressor peptide by Drude simulation (D).The mean value of 3 replicas is represented as a red line. Secondary structure elements are shown on the x axis: alpha helices (h1 – h12) as green, and beta strands (s1 – s3) as blue rectangles.

For the PPARγ - NCoR system, the differences are less prominent, the RMSF curves for both the AA and Drude simulations are similar, albeit with differences in the H9 - H10 loop and the H12 and corepressor peptide region (Fig. 6C,D). Higher variability is found in the AA simulation around H2’ and the Ω loop, and also in the loop H9 - H10. Comparing the apo and corepressor bound PPARγ systems, we see the difference in the β-sheet region and H6. For both AA and Drude simulations, adding the corepressor peptide lowered the replica-averaged RMSF. With the Drude simulations, the variability among replicas is also much smaller. In cases of stable structures, like the LBD/corepressor peptide complex, the use of the Drude model seems to provide additional stabilization of the protein complex.

The radially averaged fluctuations of all heavy atoms were calculated (see Fig. S2). As for ubiquitin, we notice the increase in the fluctuations as we move towards the residues located further from the center of geometry. The fluctuations are higher in the Drude simulations than in the AA simulations, reaching up to 3.3 Å in the apo system and 5.3 Å in the corepressor-bound system near the surface of the protein. The radius of gyration is similar for all four systems, with values of 20.1 Å (std_dev: 0.04) for AA apo system, 20.1 Å (std_dev: 0.01) for AA corepressor-bound system, 19.9 Å (std_dev: 0.15) for the drude apo system, and 20 Å (std_dev: 0.11) for the Drude corepressor-bound system.

#### Electric dipole moments from the AA and Drude simulations

Electric dipole moments of proteins contribute to the structural dynamics and function by influencing how proteins interact with their environment. Accurate modeling of the dipole moment can thus improve the overall representation of intermolecular interactions. The dipole moments of ubiquitin and PPARγ were calculated along the trajectories. The data are presented as time series of the dipole moment of the full protein. We further calculated the average dipole moment of the protein backbone by-residue.

For ubiquitin, the comparison of the dipole time series calculated from the AA and the Drude simulations shows that, overall, both force fields give average values and fluctuations with roughly the same magnitude (Fig. S3). The by-residue analysis shows that the amino acids in alpha helices systematically show larger backbone dipole moments in the Drude model than in the AA force field (Fig. S4), similar observations were made by Lopes et al ^6^. In beta sheets, the opposite is generally observed, that the backbone dipole moments calculated for the Drude simulations are lower than those in the AA simulations.

We calculated dipole moment timeseries from the AA and Drude simulations of the apo PPARγ protein and the apo PPARγ protein complexed to the co-repressor peptide (Fig 7). For the PPARγ apo system, we see lower dipole values in simulations with AA force field, with the average of 247 D, compared to the Drude force field, where the average value is 329 D.

**Figure 7.**
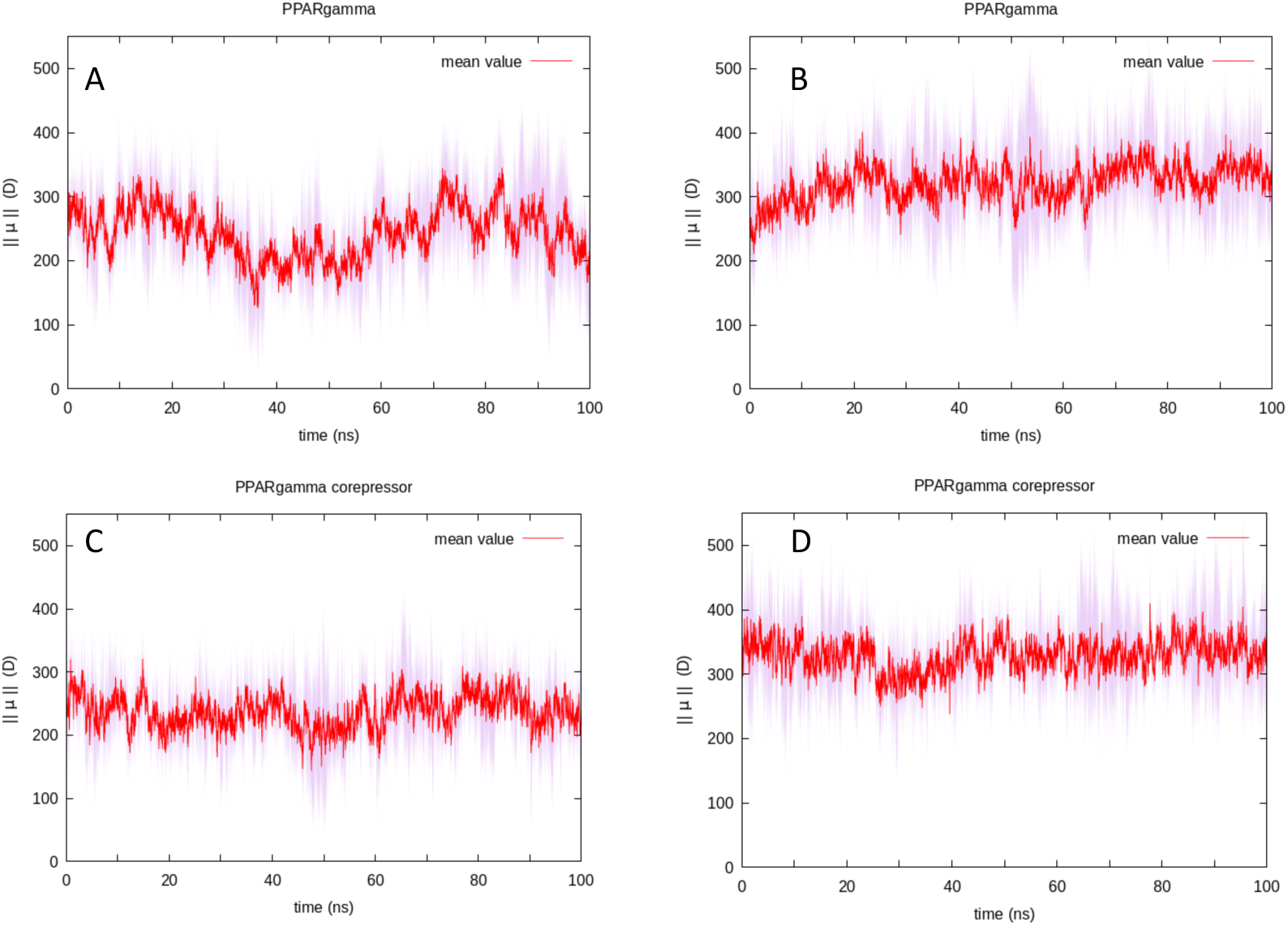
Protein dipole moment timeseries of the PPARγ LBD by AA simulation (A), PPARγ LBD by Drude simulation (B). PPARγ LBD bound to the corepressor peptide by AA simulation (C); PPARγ LBD bound to the corepressor peptide by Drude simulation (D). The mean value of 3 replicas is represented as a red line.

The calculations of PPARγ corepressor bound system follow the same trend, where the average values are 240D and 332D for the AA and the Drude FF, respectively.

Concerning the by-residue average dipole moment, we see that in the case of PPARγ, the variation of residue dipole moments is much more significant than what was observed in ubiquitin for both the apo protein and the protein complexed with the co-repressor peptide (Fig. 8). Systematically, the dipole moments in alpha helices are larger in the Drude simulations than in the AA force field simulations for PPARγ.

**Figure 8.**
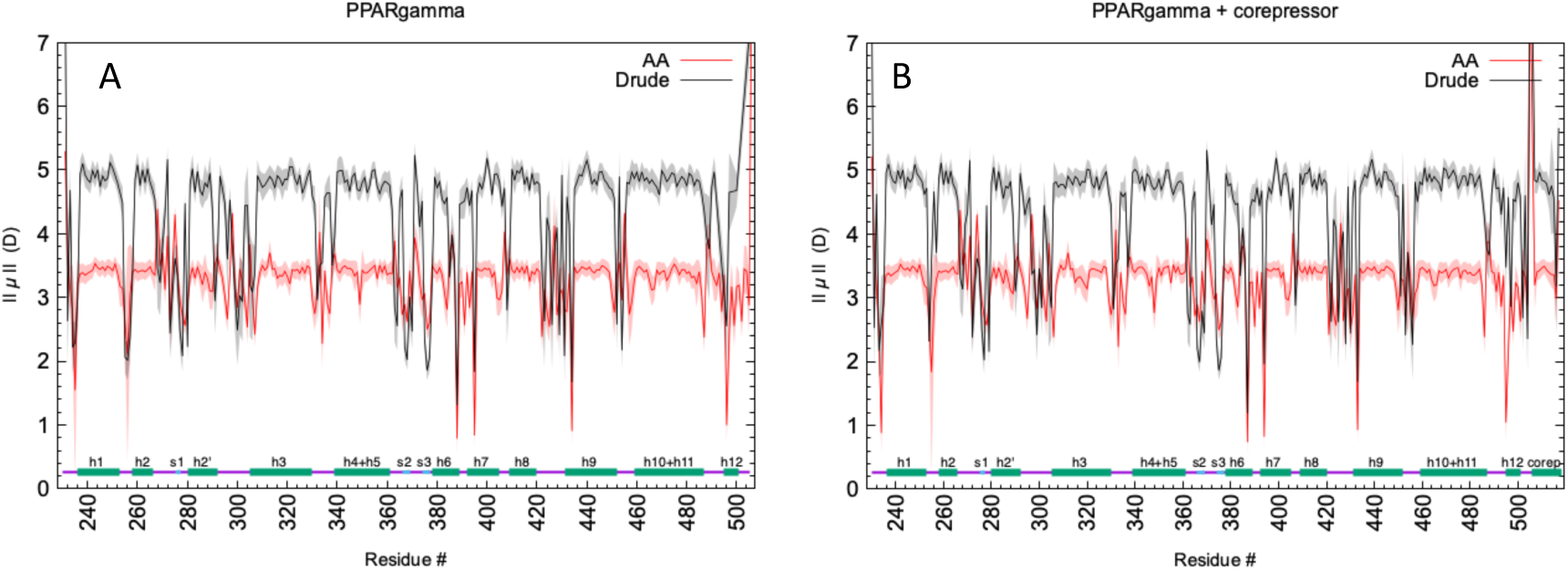
By-residue dipole moments of PPARγ for the apo protein (A) and for the LBD in complex with the co-repressor peptide (B). Secondary structure elements are shown on the x axis: alpha helices (h1 – h12) as green, and beta strands (s1 – s3) as blue rectangles.

#### Correlated motions

Correlated motions are important for understanding how the motions in different regions of the protein are coupled to other regions and how those coupled motions change in response to different perturbations, such as ligand binding. Changes in the correlated motions can effectively occur over long distances. In its simplest interpretation, this could correspond to allostery in the absence of conformational changes ^41^. It is therefore important to identify the residues involved in this transmission of structural dynamic information. This information can be obtained by calculating the cross-correlations, which complement the fluctuation analysis presented above by providing information on correlated motions as calculated by Eq. 1. From the *C_ij_* correlation coefficients, which are organized as a matrix, a cross-correlation map was calculated using a color-coded 2D representation. These calculations find use in many different applications^25,42,43^.

#### Polarization softens the correlated motions

The correlated motions were calculated from the AA and Drude molecular dynamics simulations using Eq. 1, the maps are given in Fig. 9. As these maps are normalized, the correlation values range from − 1 for perfectly anticorrelated motion to +1 for perfectly correlated motion, which is found for the self-correlations of each atom. The lower triangle corresponds to the correlation map for ubiquitin calculated from the simulation using the AA force field, while the upper triangle corresponds to the correlated motions for ubiquitin calculated from the Drude simulation. We see in the lower triangle the presence of strong correlated and anticorrelated motions involving the b strands and a helices. Particularly noteworthy are the correlated motions involved b strand 1, 2 and 5, which form the b-sheet. Correlations are also seen between b strand 5 and b strands 3 and 4. In the Drude simulations, the motions are less correlated, although the correlated motions between b1, b2 and b5 are still present. The introduction of polarization via the Drude model does not significantly change the correlation pattern, but it seems to “soften” some of the correlated motions. Interestingly, the motions of loop β1-β2 and loop α1-β3 changed character from somewhat correlated to somewhat anticorrelated motion. These regions of the protein have been reported to exhibit a pincer-like motion. However, given that the values in both cases are near zero, this suggest that the motion remains complex and not purely correlated or anti-correlated.

**Figure 9:**
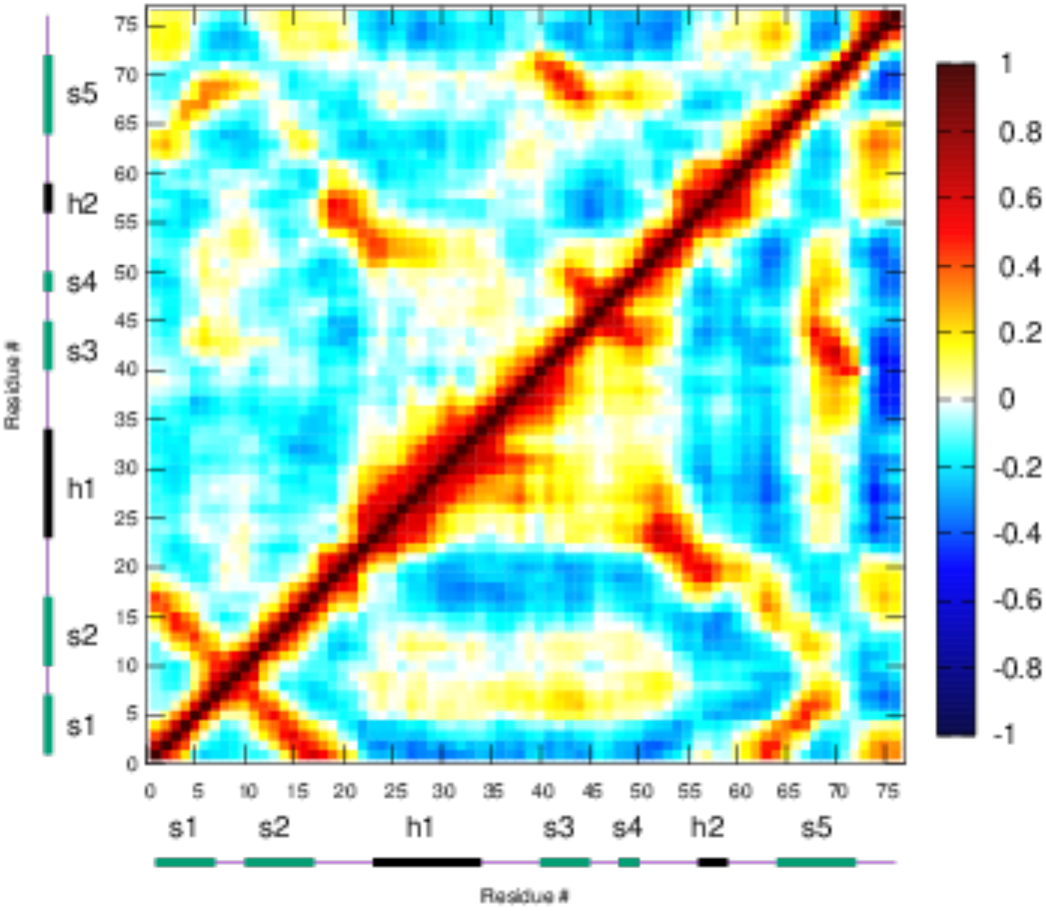
Correlated motions of ubiquitin calculated from the molecular dynamics simulations (a) from the Drude simulation (upper triangle), compared to the AA simulations, mower triangle. Correlated motion maps are represented with a color code related to the sign and intensity of correlations (ranging from dark blue for perfect anticorrelations to dark red for perfect correlations). The secondary structural elements are indicated.

We assessed the effects of polarization on the collective motions from the MD simulations of the PPARγ LBD by calculating the cross-correlations as described above for ubiquitin (Fig. 10). As for ubiquitin, the lower triangle of the corresponds to the correlation map for PPARg calculated from the simulation using the AA force field, while the upper triangle corresponds to the correlated motions of PPARg calculated from the Drude simulations. The calculations were done for both the apo and corepressor-bound forms. Regarding the general aspect of the correlation maps, for both forms, we notice a great similarity between the two. The differences are noticeable regarding the intensities of the correlations, as they appear damped in the Drude simulations, to the point that in particular regions, correlation islands disappear. In the case of the PPARγ apo system, the most significant isles represent the correlation between H12 - Ω loop residues, and H1-H9. While strongly present in the AA maps, only traces of the isles are present in the Drude simulations. Concerning the PPARγ bound to the corepressor, the most important distinction in the correlated motions is observed between helices 1 and 10, and in the motions between the H12 – H3 and H12 - H4. The peptide itself seems to be positively correlated to the C-terminal end of H3 and the N-terminal end of H4, which is a functionally important interaction, considering that the H3, H4 and H12 constitute the platform for corepressor binding.

**Figure 10.**
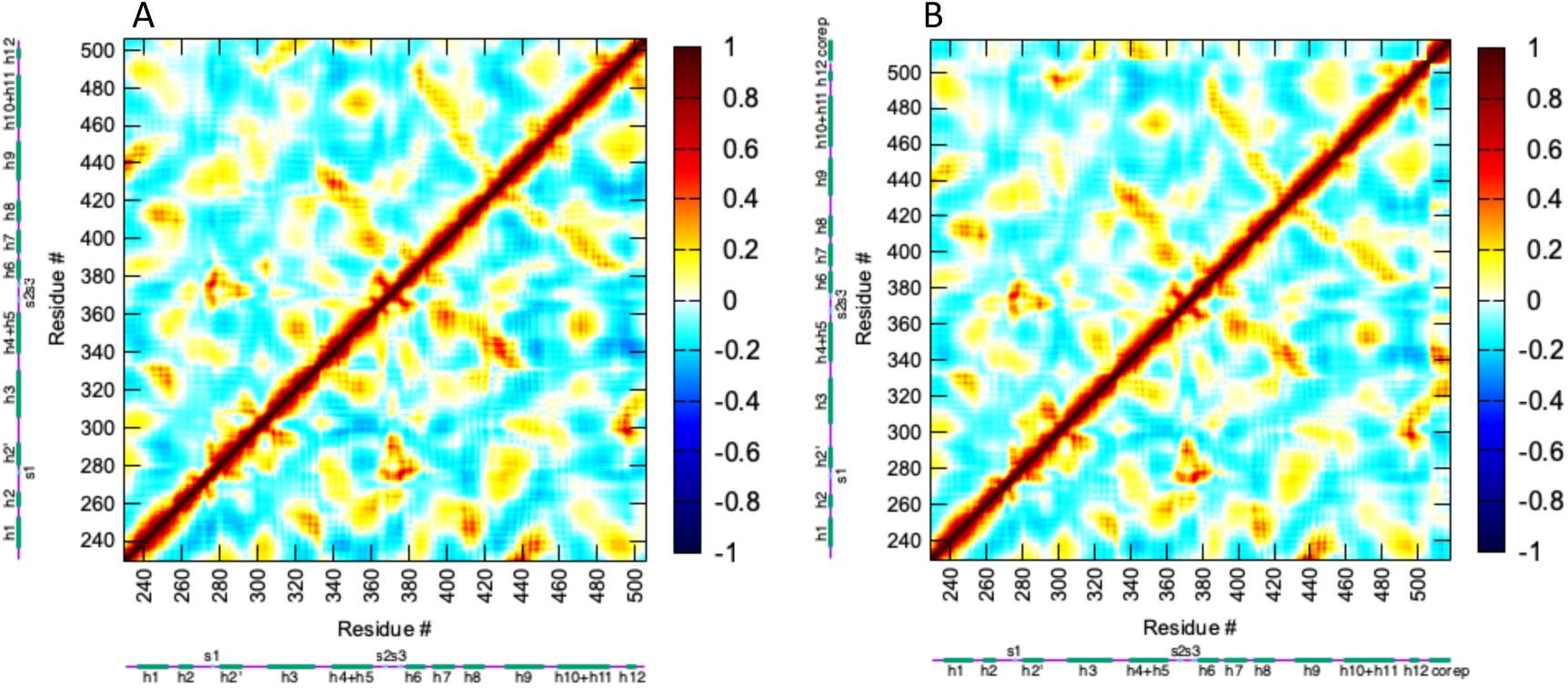
Correlated motions calculated from the simulation of the PPARγ LBD apo form (A), and corepressor bound form (B). Correlated motion maps are represented with a color code related to the sign and intensity of correlations (ranging from dark blue for perfect anticorrelations to dark red for perfect correlations). The secondary structural elements are indicated.

#### Community Network Analysis provides further interpretation of the correlated motions

To further interpret the consequences of the long-range correlated motions, we performed a community network analysis (CNA). Maps from a CNA are derived from a functional clustering of correlated motions obtained from MD simulations. It has been shown that this type of analysis, based on the Gervan-Newman algorithm, can be used to interpret long range communication and dynamic allostery of proteins ^44,45^. Community maps can help interpret how different parts of proteins move together and how changes in one part of a protein can affect the dynamics of distant sites. Communities highlight regions of the protein that exhibit collective movements and may represent functionally important domains or allosteric communication pathways. Using correlation matrices calculated from our MD simulations and the R package bio3D ^36^, we obtained coarse grained networks of dynamically coupled communities. Community network maps are depicted using colored spheres mapped onto the average 3D structure, in tube representation. The size of a node is related to the size of its community and larger spheres depict larger number of residues. The edges connect coupled communities, where thicker edges correspond to higher degree of correlation.

The results from the community network analysis of ubiquitin using the correlated motions from the AA simulation are shown in Fig. 11A. The specific compositions of the nodes are given in Supplementary Material. The protein structure is represented as a tube with the nodes, which correspond to individual residues or groups of residues, and edges, which represent dynamic correlations between nodes, are superposed. The thickness of the edges indicates the strength of the correlation. The different communities are color-coded and labeled. The analysis of the AA simulation of ubiquitin resulted in 7 community nodes and 8 edges. The most prominent node puts residues of loop S2 – H1 and residues of H2 in the same community node, N3, a large node containing 18 residues. These regions show a high degree of correlated motion between them and are therefore grouped together in a single node, see Fig. 11A. Node 3 has connections to other nodes (Ns 1, 4, 6 and 7), suggesting that this region is highly correlated to much of the protein. There is a strong edge to N4, which encompasses almost all of helix 1 and residues of the loop between helix 1 and beta strand 3. Node 4 makes a strong connection to N5, which encompasses the C-terminal end of S5 and part of S3. These connections between communities are also seen in the correlation map (Fig. 9). The correlation between H1 (N4) and S5 (N5) is not represented by one community node, but rather two nodes connected by an edge running between them. This analysis highlights the topological features of ubiquitin identifying correlation along secondary structural elements and highlighting the importance of the region loop S2-H1 and H2 for the rest of the protein. This region is also the region that has shown the pincer motion.

**Figure 11.**
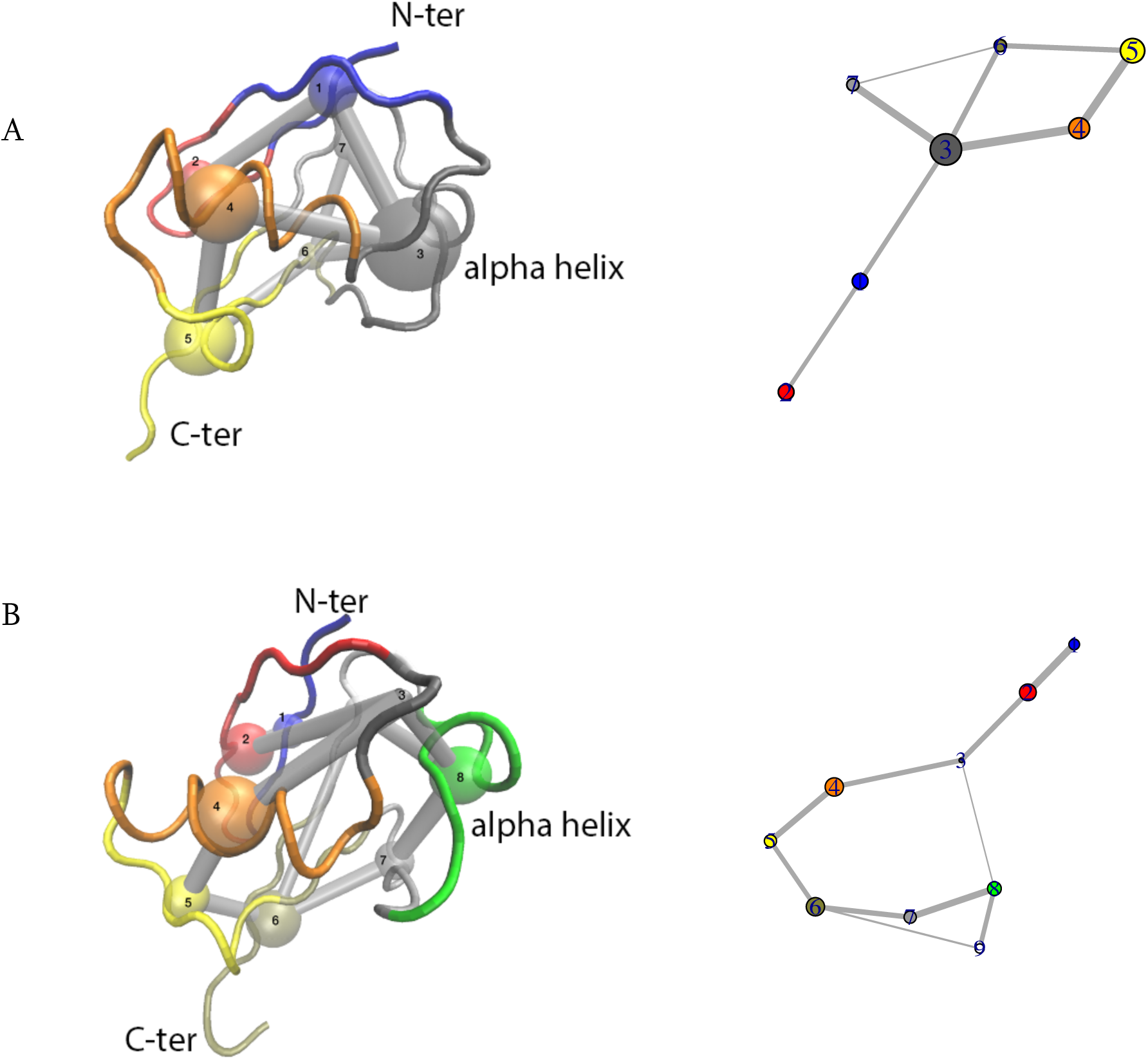
Community network analysis of Ubiquitin dynamics. On the left, the colored nodes are superposed on the protein backbone structure, represented as a tube and colored according to the nodes. The edges are denoted as grey connections between the nodes, where the thickness indicates the strength of the correlation between two nodes. On the right is the network representation. In (A) are the results from the AA simulation, and in (B) are the results from the Drude simulations.

The CNA analysis of the Drude simulation of ubiquitin gives 9 community nodes and 10 edges (Fig. 11B). The higher number of nodes suggests that there is more decoupled motion in the simulation of this protein. The nodes tend to be smaller, especially noted for N3 containing only 3 residues (residues 19:21). In the AA simulation, this region showed much stronger correlations. We see practically all of the secondary structure elements having their own nodes, with the exception beta strands S3 and S5, which have one community built around them. Since the correlations used in the community analysis are calculated from the MD simulations, that this analysis did not capture any highly correlated motions is probably due to the less intense correlated motions in the Drude simulations. The specific compositions of the nodes are given in Supplementary material.

#### CNA analysis of PPARγ

The CNA of the PPARγ apo system using the correlated motions calculated and averaged from the 3 replicas of molecular dynamics simulations detected 13 nodes for the AA simulations and 11 nodes for Drude simulations (Fig. 12).

**Figure 12.**
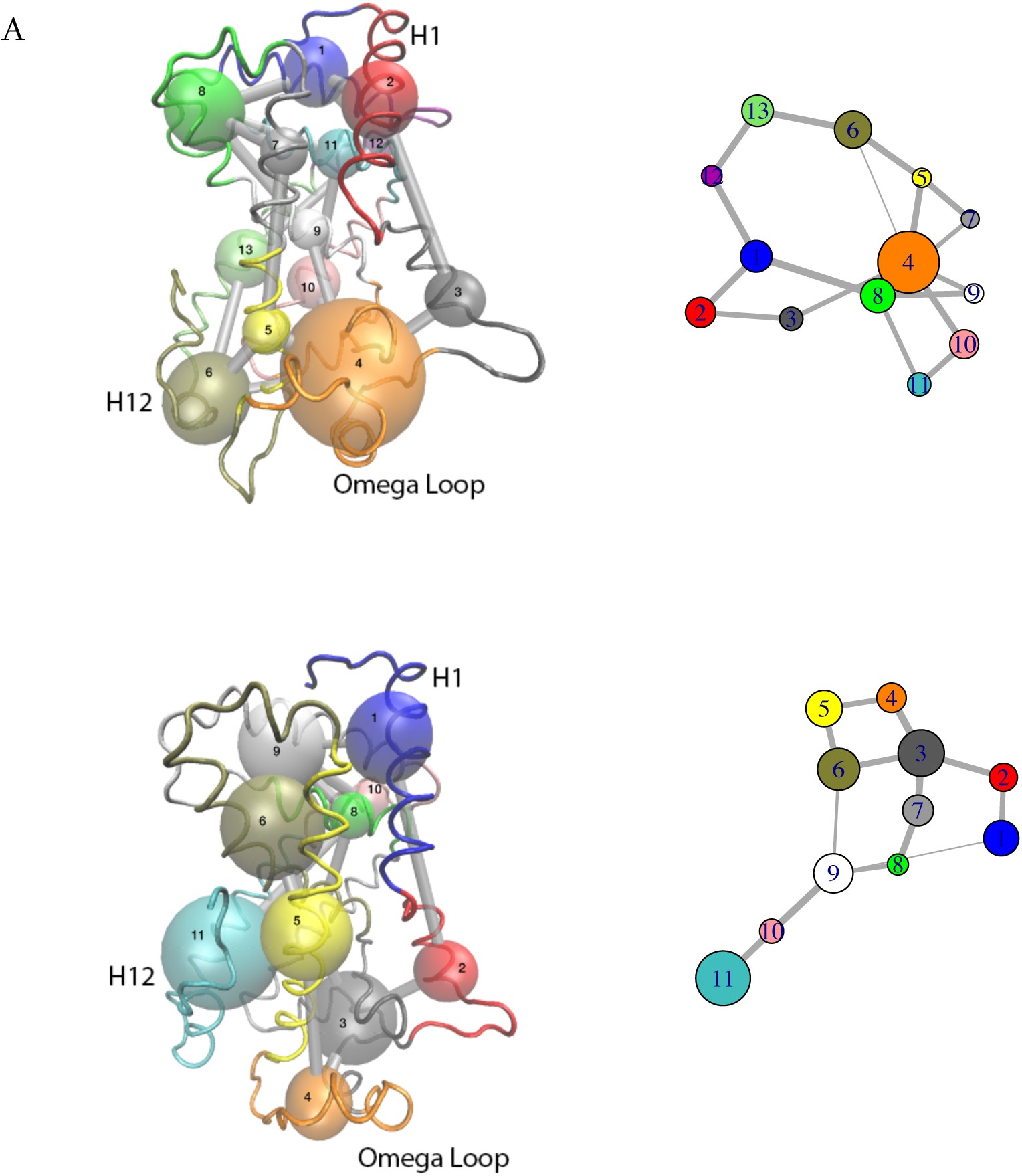
Community network analysis of the PPARγ LBD. On the left, the colored nodes are superposed on the protein backbone structure, represented as a tube and colored according to the nodes. The edges are denoted as grey connections between the nodes, where the thickness indicates the strength of the correlation between two nodes. On the right is the network representation. In (A) are the results from the AA simulation, and in (B) are the results from the Drude simulations.

The specific compositions of the nodes are given in Supplementary material. The nodes correspond largely to entire secondary structure elements, mostly helices, however, the AA simulations have four nodes encompassing spatially adjacent residues belonging to different helices: the first community regroups the N-terminal residues with residues from H9 (node 1, 23 residues, 230:234, 432:449), the second node regroups H2’, the Ω loop, the beta sheet and helix 6 (node 4, 45 residues, 274:296, 363:384), the third one regroups the Ω loop C-terminal residues with H12 residues (node 6, 27 residues, 298:306, 488:505), and the fourth one regroups the loop between H3 and H4 along with H4 with the H8 – H9 loop (node 8, 25 residues, 333:348, 423:431)). Node 4 is the one most coupled to other nodes in the apo structure. Interestingly, node 4 shows a relatively weak direct coupling to node 6, which contains the transcriptionally important H12, but it is strongly coupled to node 5, which encompasses the N-terminal end of H3. This lack of strong direct coupling may be due to the fact that the spatially near loop in node 6 is quite flexible. There is also a relatively strong coupling between the loop H8-H9 (node 8) with the rest of the protein. Interestingly, this loop is known to interact with cyclin H in other nuclear receptors, in particular RARα ^46^. The PPARγ LBD is known to interact with cyclin D ^47^ in the context of regulating adipogenesis. The nodes encompassing the loop regions at either end of Helix 9 are well connected to the rest of the protein and are known to be important in the allostery related to phosphorylation in other receptors ^46,48,49^.

The CNA of the Drude simulation of the apo PPARγ LBD generally shows smaller nodes than those observed in the AA simulation. The largest node, node 3 (residues 275:284, 362:384) encompasses the residues of H6, the β sheet and some of the Ω loop; the equivalent node in the AA simulation is node 4, however, the node from the Drude simulation is smaller. Many of the other nodes are along secondary structure elements. As in the AA simulation, there is no direct coupling between N3 and the helix 12 region of PPAR. In the Drude results, the coupling passes through 3 to 4 nodes depending on the path, while in the AA simulation, the coupling was either direct (weak) or through just one additional node. In addition to the couplings being different between the AA and the Drude simulation, these results suggest that the coupling between different regions of the PPAR ligand binding domain is less strong in the Drude simulations than in the AA simulations.

For the PPARγ/corepressor complex, the CNA identified 13 community nodes for correlated motions from both the AA and the drude simulations (Fig. 13). The specific node compositions are provided in Supplementary material. In both cases, the protein has 12 nodes and the corepressor peptide forms its own node. More of the nodes identified in the AA simulations include sequentially distant, but spatially near resides (nodes 1, 4 and 5), while in the Drude simulation, there is only one node that includes sequentially distant, but spatially near residues (node 3). In the AA system, there are three nodes which connect to neighboring nodes: the first groups N-terminal residues with H9 (node 1, 230:232, 431:455), the second groups the N-terminal of the Ω loop with the β sheet (node 4, 32 residues, 274:289, 372:378), and the third one associates the Ω loop with H12 (node 5, 38 residues 290:308, 487:505). In the Drude simulation, node 3 encompasses the β sheet and the end terminal end of the Ω loop.

**Figure 13.**
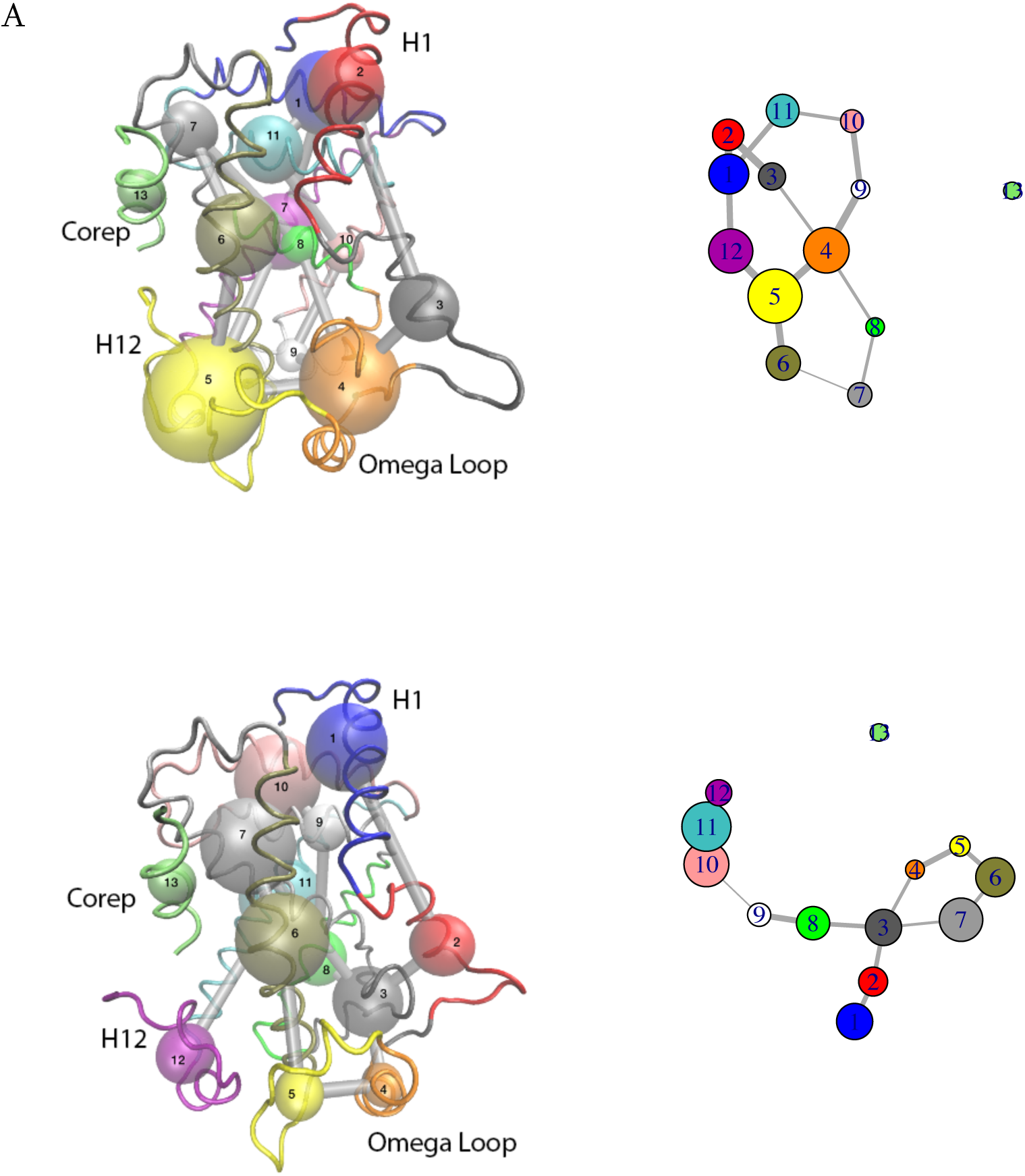
Community network analysis of PPARγ in complex with corepressor peptide. On the left, the colored nodes are superposed on the protein backbone structure, represented as a tube and colored according to the nodes. The edges are denoted as grey connections between the nodes, where the thickness indicates the strength of the correlation between two nodes. On the right is the network representation. Representation on the PPARγ LBD corepressor bound form, from AA simulation (A), and Drude simulation (B).

The CNA shows that many of the node interconnections are along the secondary structure elements, but the AA simulations display several edges between nodes beyond secondary structure. The node organization for AA is more global, as opposed to the Drude simulation results, where there is less interconnectivity and the network is more extended. Central to the interconnectivity in PPARγ is the node that encompasses the β sheet and part of the Ω loop region. This node forms a hub through which many edges connect.

CNA analysis for both the AA and the Drude simulations show the corepressor peptide as a single node. And, we further notice that this node does not form any edges to any nodes in the ligand binding domain of PPARγ. If we look back at the correlation plots of AA simulations, we see positive correlation between the corepressor peptide residues and the N-terminal of H4, while the Drude simulations did not capture these correlations. The correlations were weak and, as they did do not go over the 0.5 threshold, they are not represented by an edge. The addition of the corepressor peptide in the AA and Drude simulations does not seem to break the community of the H3 – 4 loop and H4 residues, which forms the corepressor binding platform. In the AA results, the presence of the peptide seems to increase the H12 and Ω loop community, passing from 27 (node 6) to 38 (node 5) residues, and reinforcing their correlations. In the Drude simulations, the addition of the peptide seems to decouple two different communities. First, the H11 – H12 community is split into two separate ones, connected by an edge (from node 11 to nodes 11 and 12). The second community, built around the Ω loop (node 4), is divided into 2 separate nodes (nodes 4 and 5), connected by edges. Other nodes do not seem to be affected by the corepressor addition.

One significant distinction between the simulations using the two different force fields concerns the community that represents helix H12. In the AA simulations of both apo and corepressor-bound systems, H12 and one part of the Ω loop are coupled and are therefore represented by one community. This node is of medium size, with 27 residues for the apo form and with 38 residues for the corepressor-bound form. In the Drude simulations, H12, together with H11 make up an individual community, represented by a node containing 39 residues. This community is decoupled from the node encompassing the Ω loop in both systems simulated by the Drude FF meaning there are no edge connections between them. This suggests that the correlations in the Drude simulations are not sufficiently strong to result in the CNA analysis detecting direct communication between them. Interestingly, two mutations in the omega loop region have been identified as being activating mutations in luminal bladder cancer ^50^.

In the corepressor-bound form, H12 is further decoupled from the H11, having its own community of helix residues connected by an edge to the H11. Furthermore, in both systems simulated with AA FF, the N-terminal residues of PPARγ are grouped in the same community with H9 residues, while in the Drude simulations, these N-terminal residues are in the same community as H1 residues. This coincides with high RMSF values for the N-terminal residues of the LBD in both Drude simulated systems, Fig. 6.

Helix H12 represents the activation function 2 in LBDs and therefore is physiologically important for the regulation of PPARγ’s transcriptional activity. In the transcriptionally inactive form studied here (apo or corepressor bound forms), the H12 exhibits higher flexibility and is capable of exploring multiple conformations ^51^. In the community analysis of the Drude simulations, we noticed the decoupling of the H12 from the other regions, notably the Ω loop and the H11. This suggests that these regions explore different movements which are not directly correlated and display different conformational dynamics. The lack of high correlating communities and the presence of communities largely representative of individual alpha-helices is apparent in the correlation maps, where the Drude simulations display attenuated colors, and thus smaller correlations. It is generally appreciated that around the ligand binding pocket of the PPARγ LBD, the region is flexible in the absence of a ligand, so we would expect a low degree of correlated motion is this area, notably of the functionally relevant helix H12 and the conformationally flexible Ω loop.

#### Shortest Path Method (SPM)

A second approach for interpreting correlated motions is the Shortest Path Method (SPM), which was used through the online webserver ^38^. This tool was used to assess the importance of individual residues, and their pairwise connections, in the structural dynamics of the two proteins. This is in contrast to the community network analysis, which establishes communities around multiple residues. The SPM method produces a network graph based on mean distances and correlation values. The shortest path lengths were calculated using the Dijkstra algorithm ^38^. The shortest path is the most direct path following the most significant connection between two residues and shows how the residues are connected in the structural dynamics of the protein. The tool is mostly aimed at exploring key residues implicated in enzymatic activity, but here we use it to assess the similarities and differences of simulations using different force fields.

Concerning ubiquitin, the SPM analysis using the correlated motions calculated from the AA and Drude simulations show a similar pattern (Fig 14). Interestingly, in both cases, there is a path detected between the beginning of loop β1-β2 and the loop after the C-terminus of the α-helix. These regions of ubiquitin have been identified in other works as exhibiting a pincer-type motion. For this small, tightly packed protein, there do not seem to be any noticeably difference between the AA and Drude simulations in terms of the SMPs.

**Fig 14.**
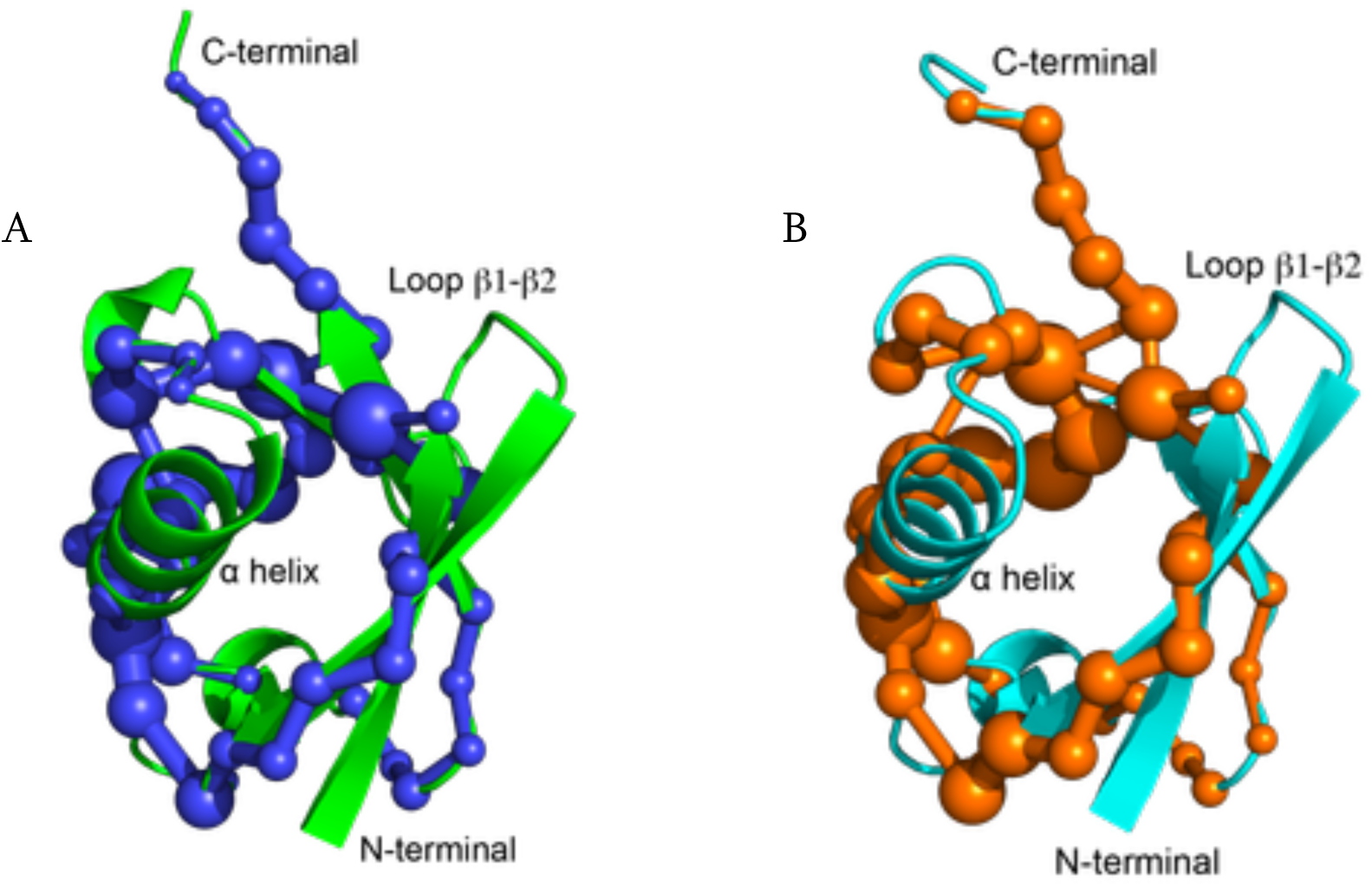
Shortest Path Method ball and stick representation mapped on ubiquitin average structure. The AA simulation result is in blue (A) and the Drude simulation result is in orange (B).

The PPARγ apo system (Fig 15) shows a graph network connecting different nodes corresponding to same secondary structure elements. For example, if we look at the side view of the structure from AA simulation, we can see a path spanning along the entire helix H10-11, and then continuing connecting the loop and H12, and even further, the Ω loop. This suggests a correlation and coupling of these secondary structure elements. On the other hand, the Drude simulation shows no such connection and the functionally important H12 is not coupled to Ω loop movements. Similar observations were made from the community network analysis.

**Figure 15.**
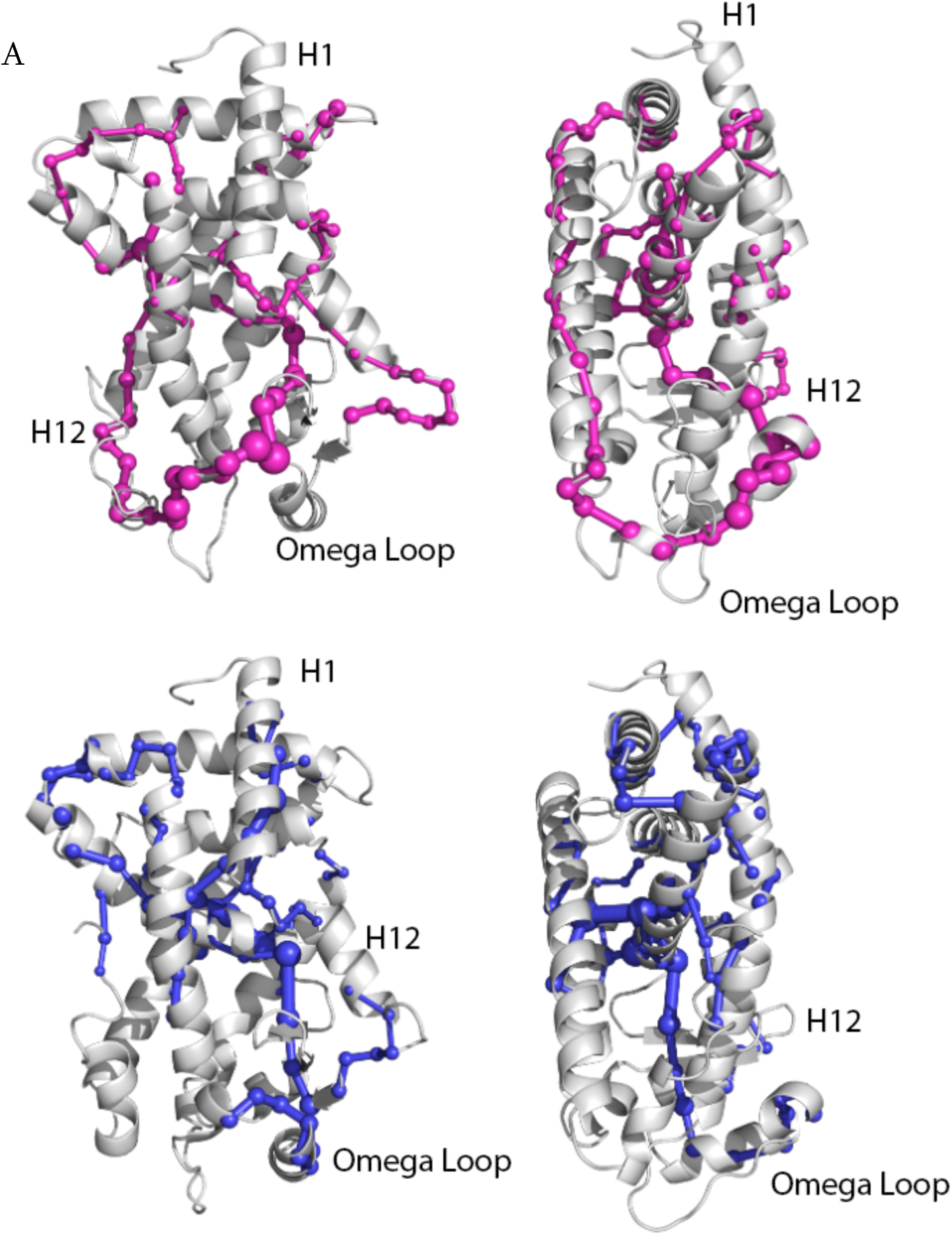
Shortest Path Method ball and stick representation mapped on the front and side views of PPARγ LBD apo form. The AA simulation result is in magenta (A) and the Drude simulation result in blue (B).

Compared to the apo PPARγ, the SPM paths of PPARγ bound to the corepressor peptide are relatively different for both the AA and Drude calculations (Fig. 16). In this case, we again discern in the case of the AA simulation, the SPM spanning throughout the ‘upper’ region of the LBD and the one in the ‘bottom’ region with respect to the illustration, where H12 and the Ω loop are connected. Interestingly, we see short paths between alpha helices.

**Figure 16.**
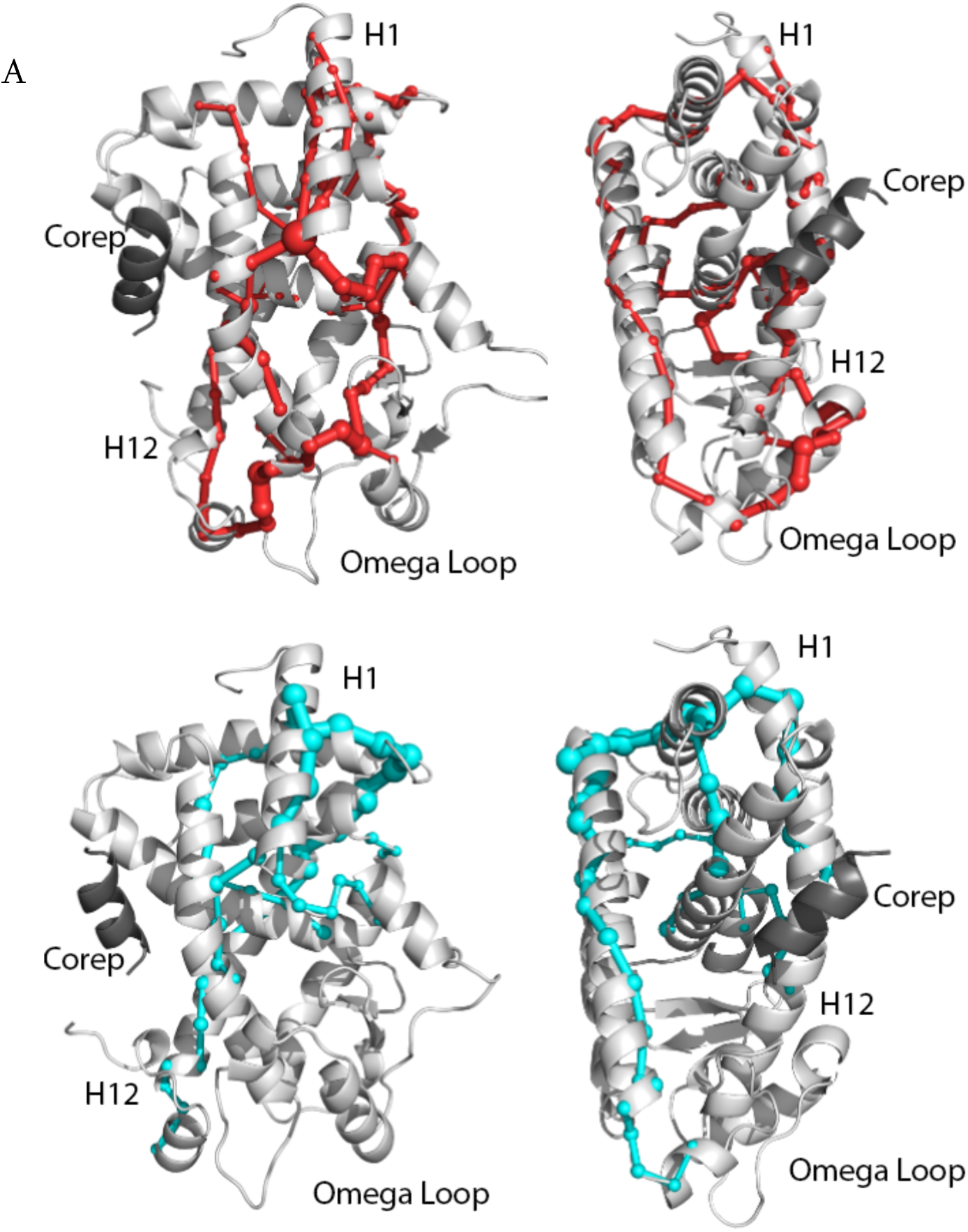
Shortest Path Method ball and stick representation mapped on the front and side views of PPARγ LBD with corepressor peptide bound. The AA simulation result is in red (A) and the Drude simulation result in cyan (B).

The use of the Drude force field leads to a decoupling of H12 and the Ω loop region in the apo protein; the same observation was made from the community network analysis. The corepressor peptide, even though it was included in the SPM network calculation, does not appear to participate in the shortest path representation. Despite the somewhat correlated motions between the corepressor peptide and helices H3 and H4 (Fig. 10B), the co-repressor peptide is not connected to the rest of the protein in this analysis. A similar conclusion was made from the community network analysis, that is, the corepressor peptide does not enter into any communication network. We also notice the absence of the SPM path in the regions of the loop H3 – H4, probably caused by the addition of the corepressor peptide. This suggests that the presence of the peptide, while not directly implicated in a network, will perturb the underlying communication network of PPARγ.

## Conclusions

In this work, we used the Drude polarizable force field in molecular dynamics simulations of two proteins, human ubiquitin and the ligand binding domain of human PPARγ. We compared the results to simulations using the CHARMM all atom atomic force field. We examined the effect of the Drude force field on standard measures of structural dynamics, such as RMSD and RMSF via comparison to simulations using a classical, AA force field. We generally found conformational changes leading to a higher RMSD and, in flexible regions of the proteins, greater flexibility. But overall, the trends remained the same. Looking at dipole moments, we confirmed what was found in other studies, that, for the most part, the Drude model leads to greater values of dipole moments for individual residues.

We also characterized for the first time the effects of using the Drude force field on correlated motions, which have been implicated in the biological function of proteins. The correlated motions were characterized by correlation maps generated by molecular dynamic simulations, by community network analysis (CNA) and Shortest Path Method (SMP) analysis. The latter two interpret information from the correlated motions obtained from the simulations. The CNA distinguished regions of the proteins where residues interact strongly with each other, and are placed in the same community, from those that interact more weakly. The latter are placed outside of the community and if they are part of another community with sufficiently strong correlations to another community, the information is indicated by connection between the nodes. The analysis further reveals paths through which signals, structural or through interactions with other proteins, can propagate from one region to another. The SPM approach provide more residue-to-residue mapping of the correlated motions, but both approaches provided insights into how protein dynamics map onto the modular organization of the protein and reveal residues or communities of residues that display coordinated motions. Such coordinated motions may underpin allosteric communication. The flexibility of certain regions and the stability of others have an impact on the outcome of this analysis, so we investigated the effects of introducing polarization into the molecular dynamics simulations through the use of the Drude force field.

Analysis of the correlated motions through examination of the correlation maps, the CNA and the SMP analysis based on these correlations show that the use of polarization via the Drude force field affects the correlated motions by decoupling the motions and generally softening the motions. Correlated motions are generally considered to arise from low frequency collective motions. This suggests that, perhaps the simulated protein will be less responsive to perturbations introduced, for example, by ligand binding or by the introduction of point mutations. So, perhaps there is an advantage to using the AA force field if one is interested in the studying allosteric behavior of proteins based on their collective motions. In conclusion, we notice that the major difference arises in regions of the protein that are known from simulations to exhibit more significant flexibility that in other regions of the protein, for example, in the ligand binding domain of PPARγ, around the ligand binding pocket, and aforementioned helix H12.

## ASSOCIATED CONTENT

### Data Availability

Initial conformations, input parameter and topology files, input configuration files for the dynamics simulations and analysis, representative output trajectories and well as datasets from analysis for each system are available by contacting the corresponding author. The NAMD and CHARMM molecular dynamics software are freely available from their respective distribution sites.

### Supporting Information

A file (pdf) containing additional figures and tables for

- Radially averaged RMSF for ubiquitin and PPARγ
- Time series for ubiquitin dipole moment
- By-residue backbone dipole moment for ubiquitin
- Timeseries for the D411-H453 salt bridge for apo-PPAR and PPARγ-complex
- Table listing the Composition of the nodes from the community network analysis of ubiquitin
- Table listing the Composition of the nodes from the community network analysis of PPARγ

## AUTHOR INFORMATION

### Author Contributions

The computer studies were conceived and executed by AM and RHS. All authors contributed to the analysis and the manuscript was written with contributions from all authors. All authors approved the final version of the manuscript.

### Notes

The authors declare no competing financial interest.

## Acknowledgements

The authors acknowledge the Agence Nationale pour la Recherche (FRIDaY project ANR20-CE44-0004) for financial support. The computational work was supported by the Centre National de la Recherche Scientifique, the Institut National de la Santé et de la Recherche Médicale and the University of Strasbourg. The authors acknowledge particularly the support of the Strasbourg University High Performance Computing Center and GENCI (Grand Equipement National de Calcul Intensif) for computing resources. AM would like to thank Dr. Nastazia Lesgidou of the Biomedical Research Foundation Academy of Athens, Greece, for discussions on Community Network Analysis. Clara C. Stote is acknowledged for help with the CNA illustrations.

## ABBREVIATIONS

LBD: Ligand binding domain.

## TOC Graphic

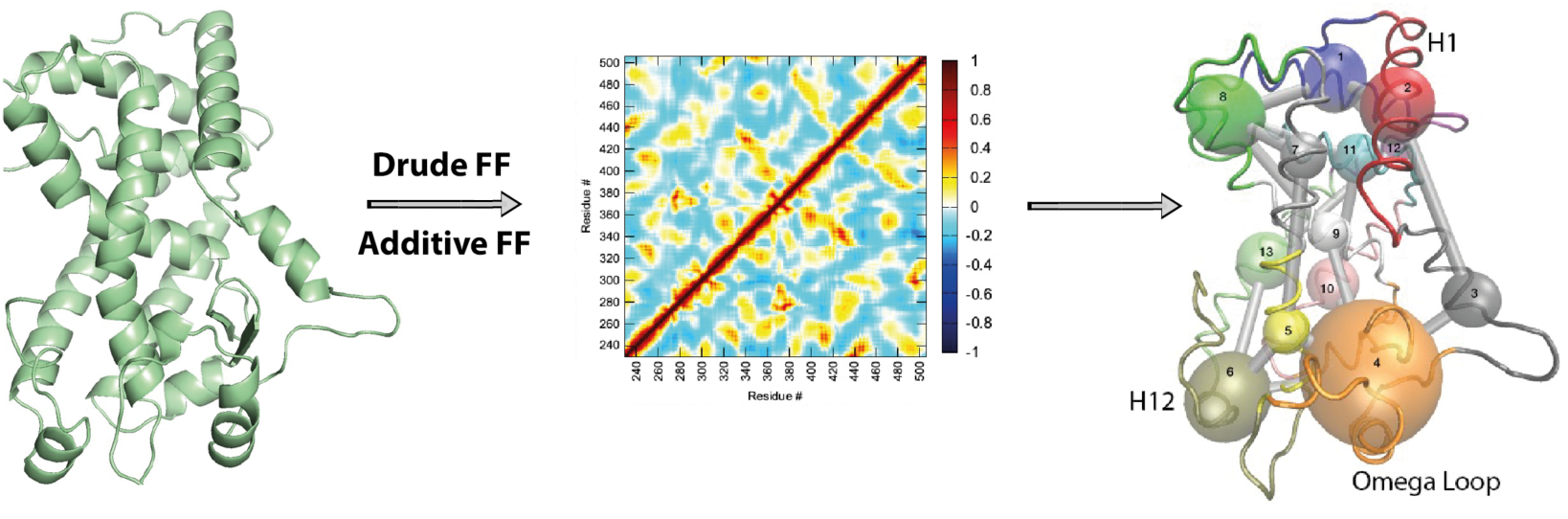

