## Supplementary Material for "Impact of force field polarization on the collective motions of proteins"

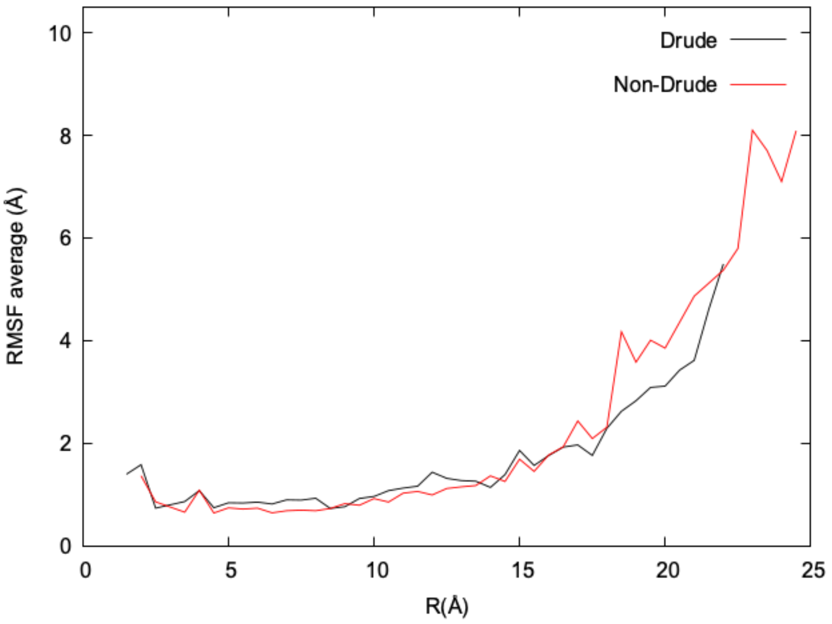

Figure S1. Radially averaged fluctuations for ubiquitin from the AA and Drude simulations.

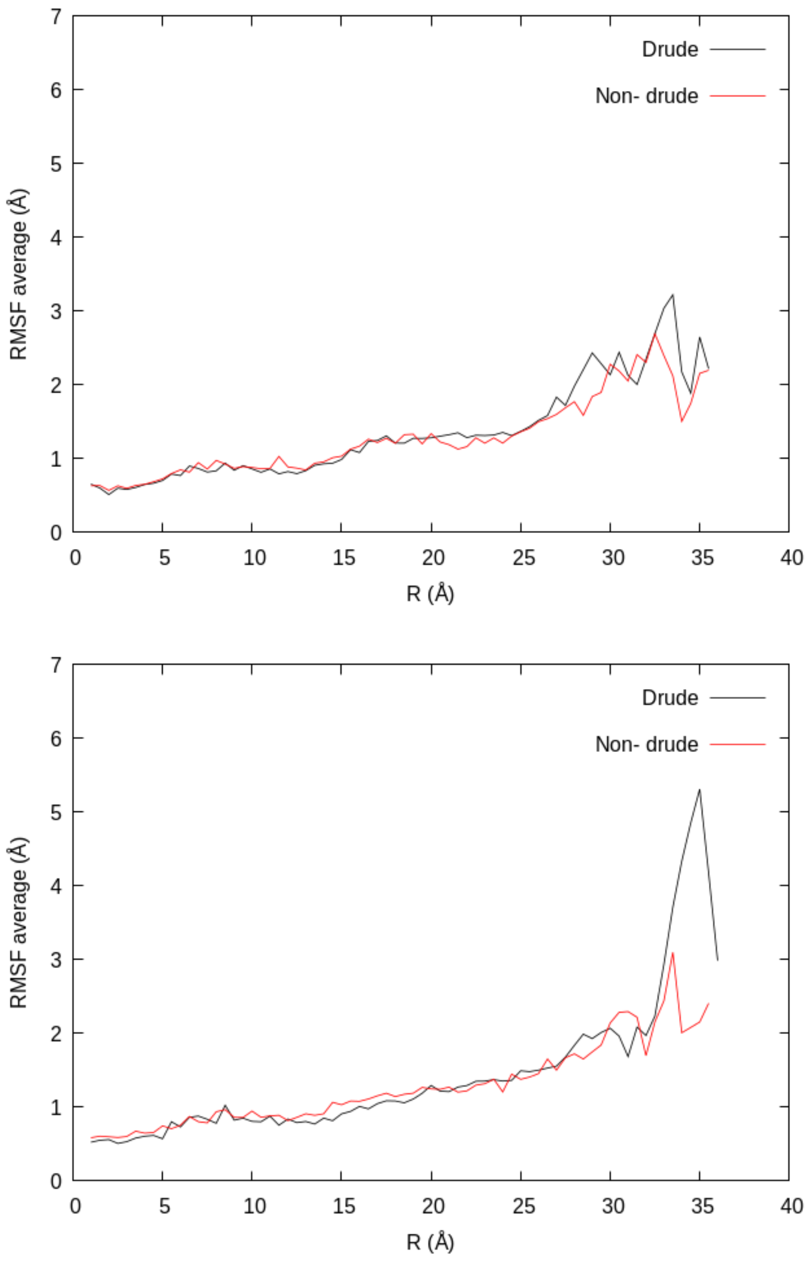

Figure S2. Radially averaged atomic fluctuations of the PPARγ LBD apo form (top), and corepressor bound form (bottom).

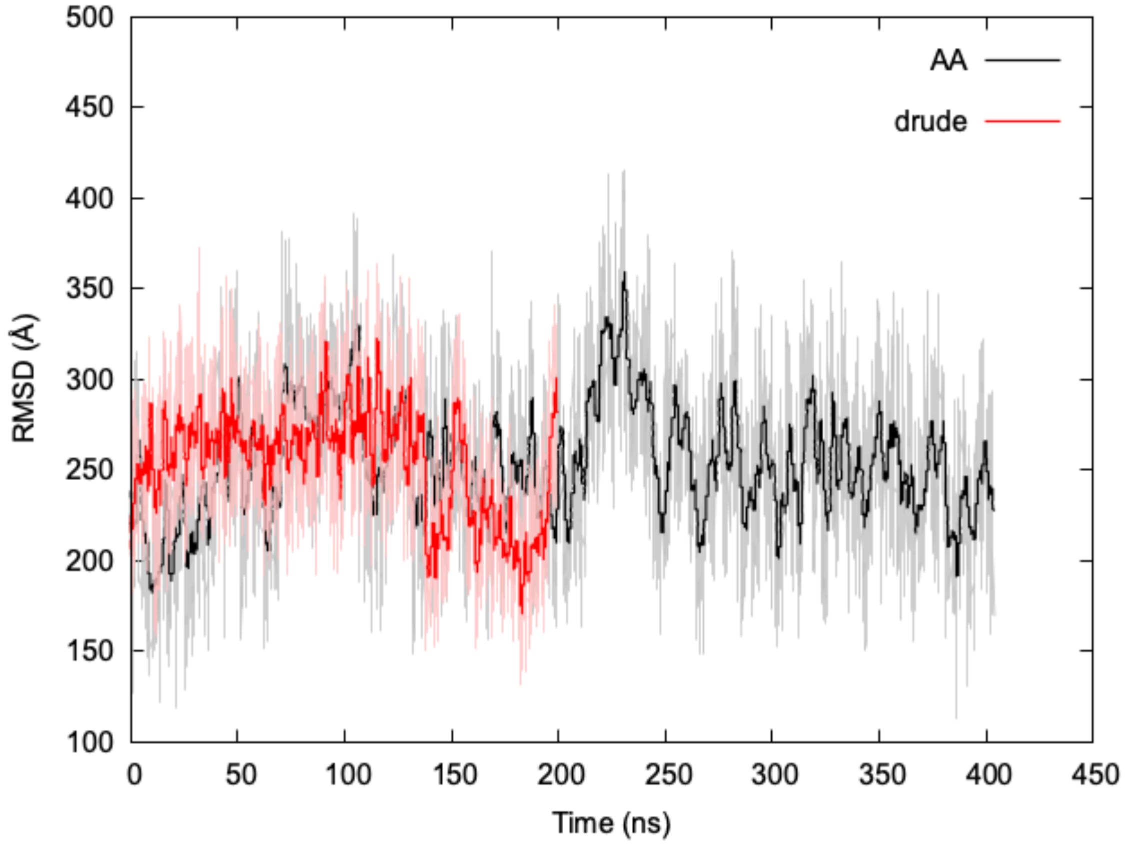

Figure S3. Time series for full dipole of ubiquitin. Shown is the running average over 100 frames in black (AA) and red (Drude) along with shades of individual values during the time series.

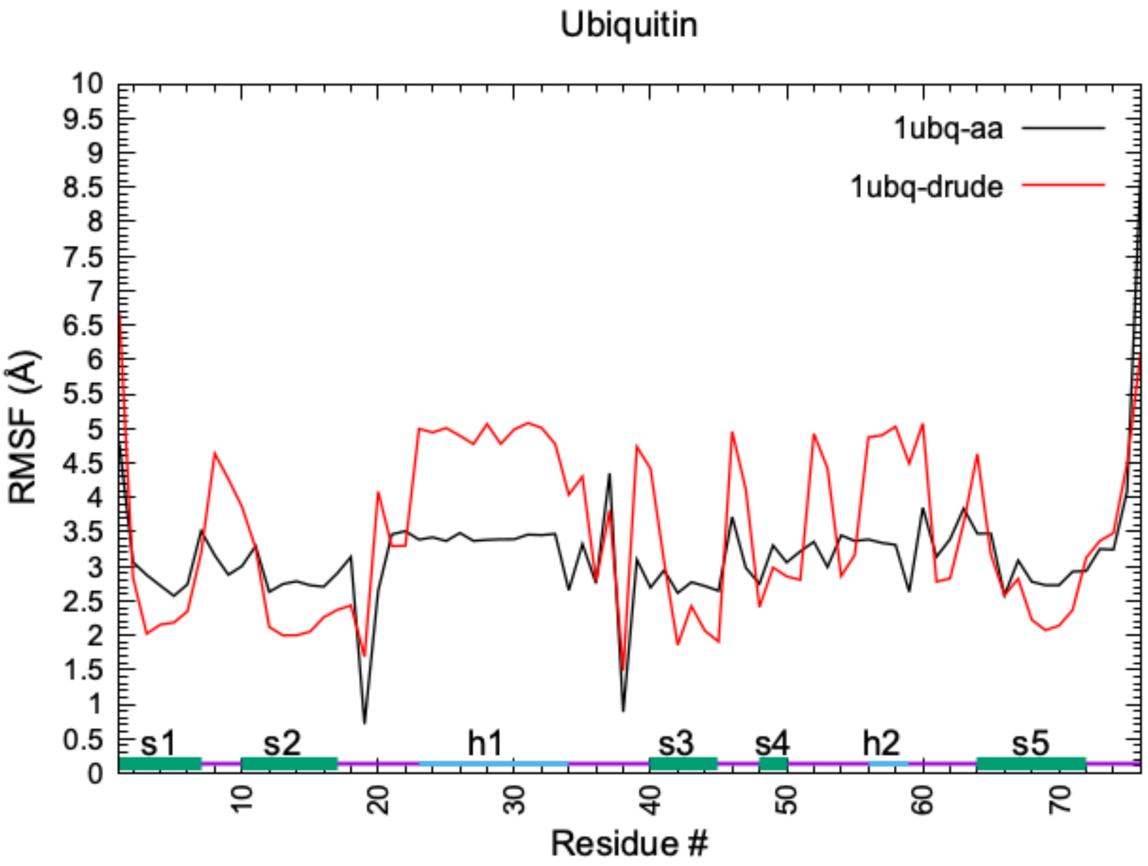

Figure S4. Average dipole moment by-residue.

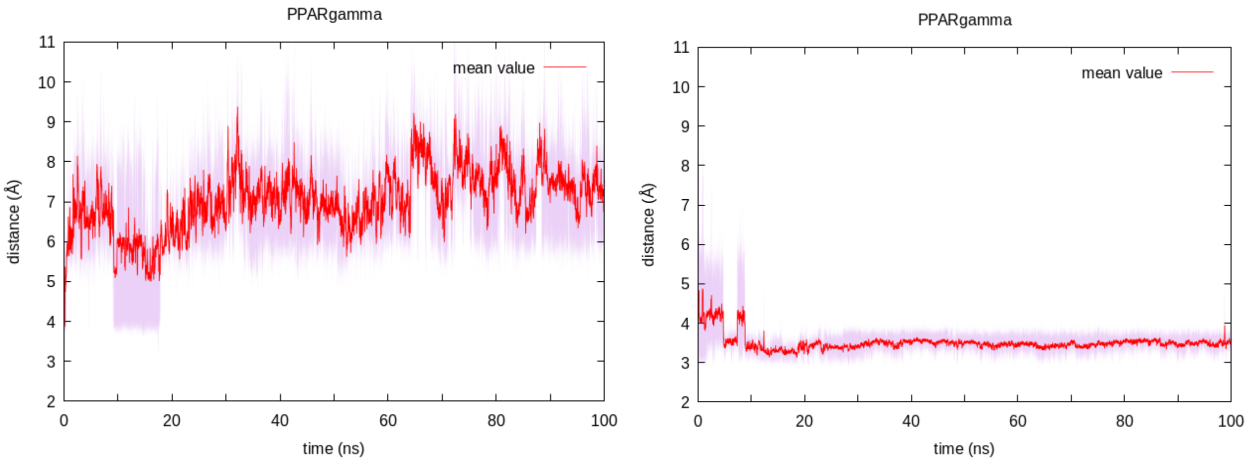

Figure S5a. Timeseries for the salt-bridge interatomic distance of the PPARγ LBD apo form, in the AA (left) and Drude simulations (right). The distance is calculated between two heavy atoms: CG of D411 residue, and NE2 of H453 residue.

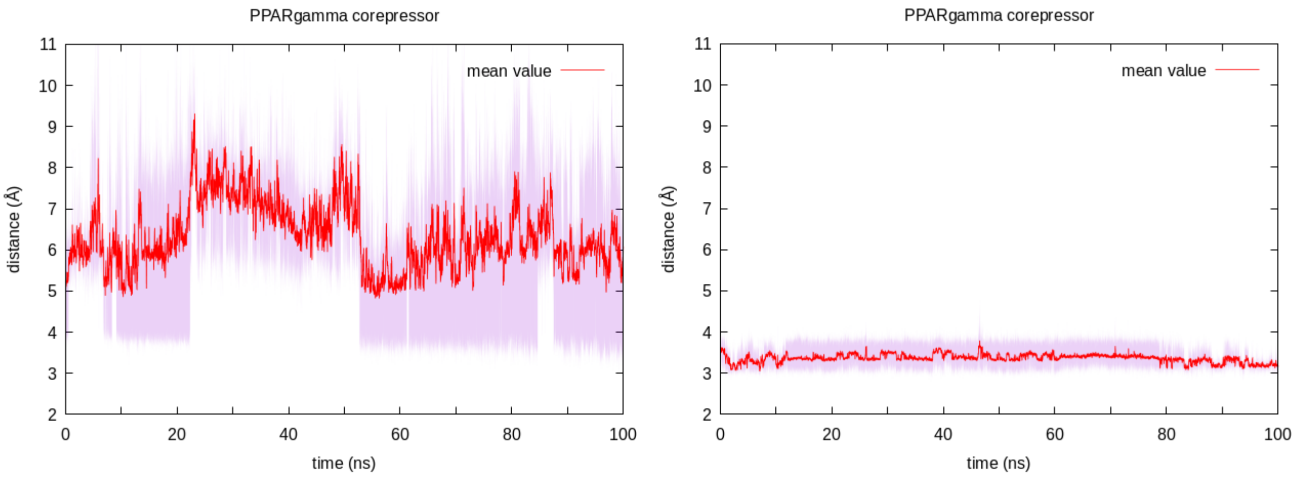

Figure S5b. Timeseries for the salt-bridge interatomic distance of the PPARγ LBD corepressor-bound form, in the AA (left) and Drude simulations (right). The distance is calculated between two heavy atoms: CG of residue D411, and NE2 of residue H453.

Table S1. Composition of the nodes from the community network analysis *of ubiquitin*. Size represents the number of amino acids in a node. Node members are the numbers of amino acids belonging to a node.

| **Ubiquitin_AA** | | | | **Ubiquitin_Drude** | | |
|---|---|---|---|---|---|---|
| node id | size | members | | node id | size | members |
| 1 | 9 | c(1 :4, 14 :18) | | 1 | 7 | 1 :7 |
| 2 | 9 | 5 :13 | | 2 | 11 | 8 :18 |
| 3 | 18 | c(19:24, 49:60) | | 3 | 3 | 19 :21 |
| 4 | 12 | 25 :36 | | 4 | 12 | 22 :33 |
| 5 | 14 | c(37:42, 69:76) | | 5 | 8 | 34 :41 |
| 6 | 7 | c(43:48, 68) | | 6 | 12 | c(42:44, 68:76) |
| 7 | 7 | 61 :67 | | 7 | 8 | 45 :52 |

Table S2. Composition of the nodes from the community network analysis. Isoform PPARγ2 numbering of LBD residues: 230 - 505. Corepressor peptide is numbered from 506 - 517. Size represents the number of amino acids in a node. Node members are the numbers of amino acids belonging to a node.

| PPAR_AA_apo | | | | PPAR_AA_corep | | |
|---|---|---|---|---|---|---|
| node id | size | members | | node id | size | members |
| 1 | 23 | c(230:234, 432:449) | | 1 | 28 | c(230:232, 431:455) |
| 2 | 22 | 235:256 | | 2 | 22 | 233:254 |
| 3 | 17 | 257:273 | | 3 | 19 | 255:273 |
| 4 | 45 | c(274:296, 363:384) | | 4 | 32 | c(274:289, 372:378) |
| 5 | 14 | c(297, 307:319) | | 5 | 38 | c(290:308, 487:505) |
| 6 | 27 | c(298:306, 488:505) | | 6 | 24 | 309:332 |
| 7 | 13 | 320:332 | | 7 | 17 | 333:349 |
| 8 | 25 | c(333:348, 423:431) | | 8 | 13 | 350:362 |
| 9 | 14 | 349:362 | | 9 | 13 | 379:391 |
| 10 | 21 | 385:405 | | 10 | 16 | 392:407 |
| 11 | 17 | 406:422 | | 11 | 23 | 408:430 |
| 12 | 15 | 450:464 | | 12 | 31 | 456:486 |
| 13 | 23 | 465:487 | | 13 | 12 | 506:517 |
| PPAR_Drude_apo | | | | PPAR_Drude_corep | | |
| node id | size | members | | node id | size | members |
| 1 | 25 | 230:254 | | 1 | 25 | 230:254 |
| 2 | 20 | 255:274 | | 2 | 20 | 255:274 |
| 3 | 33 | c(275:284, 362:384) | | 3 | 25 | c(275:277, 362:383) |
| 4 | 21 | 285:305 | | 4 | 13 | 278:290 |
| 5 | 26 | 306:331 | | 5 | 14 | 291:304 |
| 6 | 30 | 332:361 | | 6 | 27 | 305:331 |
| 7 | 22 | 385:406 | | 7 | 30 | 332:361 |
| 8 | 15 | 407:421 | | 8 | 23 | 384:406 |
| 9 | 28 | 422:449 | | 9 | 16 | 407:422 |
| 10 | 17 | 450:502 | | 10 | 31 | 423:453 |
| 11 | 39 | 467:505 | | 11 | 34 | 454:487 |
| | | | | 12 | 18 | 488:505 |
